# Cerebellar-driven cortical dynamics enable task acquisition, switching and consolidation

**DOI:** 10.1101/2022.11.14.516257

**Authors:** Joseph Pemberton, Paul Chadderton, Rui Ponte Costa

## Abstract

To drive behavior, the cortex must bridge sensory cues with future outcomes. However, the principles by which cortical networks learn such sensory-behavioural transformations remain largely elusive. Here, we posit that the cerebellum assumes a crucial role in driving cortical dynamics, thereby enabling rapid and flexible task acquisition. We introduce a computational model of cerebellar networks which learn to drive cortical networks with task-outcome predictions. First, using sensorimotor tasks we show that cerebellar feedback in the presence of minimal cortical plasticity is sufficient for rapid task acquisition and multiple task switching. Next, we demonstrate that, when trained in working memory tasks, the cerebellum can also underlie the maintenance of cognitive-specific dynamics, explaining a range of optogenetic and behavioural observations. Finally, using our model we introduce a systems consolidation theory in which task information is gradually transferred from the cerebellum to the cortex. In summary, our findings suggest that cortico-cerebellar loops play a pivotal role in task acquisition, switching, and consolidation within the brain.

## Introduction

Learning to interact with the environment requires a continuous integration of fast-changing sensory cues with future behavioural outcomes. Growing evidence suggests that cortical dynamics integrate the task-specific information that is needed for such sensory-behavioural transformations ^1–5^. One dominating view in the field assumes that cortical networks are themselves learnt or optimised leading to the rich dynamics required for task performance ^6–8^. However, to help ensure a stable representation of the world, cortical plasticity must be kept under control and relatively weak ^9–12^. This raises the question of how can the brain quickly acquire new task-specific dynamics in the presence of relatively fixed cortical connectivity.

One possible solution is to consider feedback loops that drive cortical dynamics ^13^. Computational studies have extended recurrent neural networks (RNNs) models of cortical networks (Fig. 1A) to incorporate feedback loops for task acquisition. One type of feedback loop drives RNN dynamics by projecting the readout back to the RNN ^14–16^ (Fig. 1B). Building on this line of work, two recent theoretical studies have suggested that thalamo-cortical feedback can both prepare and control RNN dynamics to achieve flexible motor sequencing ^17,18^. All of these studies assume that connectivity within the RNN itself remains fixed, thereby avoiding complex learning rules while being able to reuse RNN dynamics for different contexts ^19^. However, these approaches either assume a relatively simple feedback (i.e. a linear combination of RNN activity) or rely on theoretically optimal, but biologically implausible, derivations for the feedback signal. In particular, the possible role of more powerful, highly adaptable brain regions is often overlooked.

**Figure 1.**
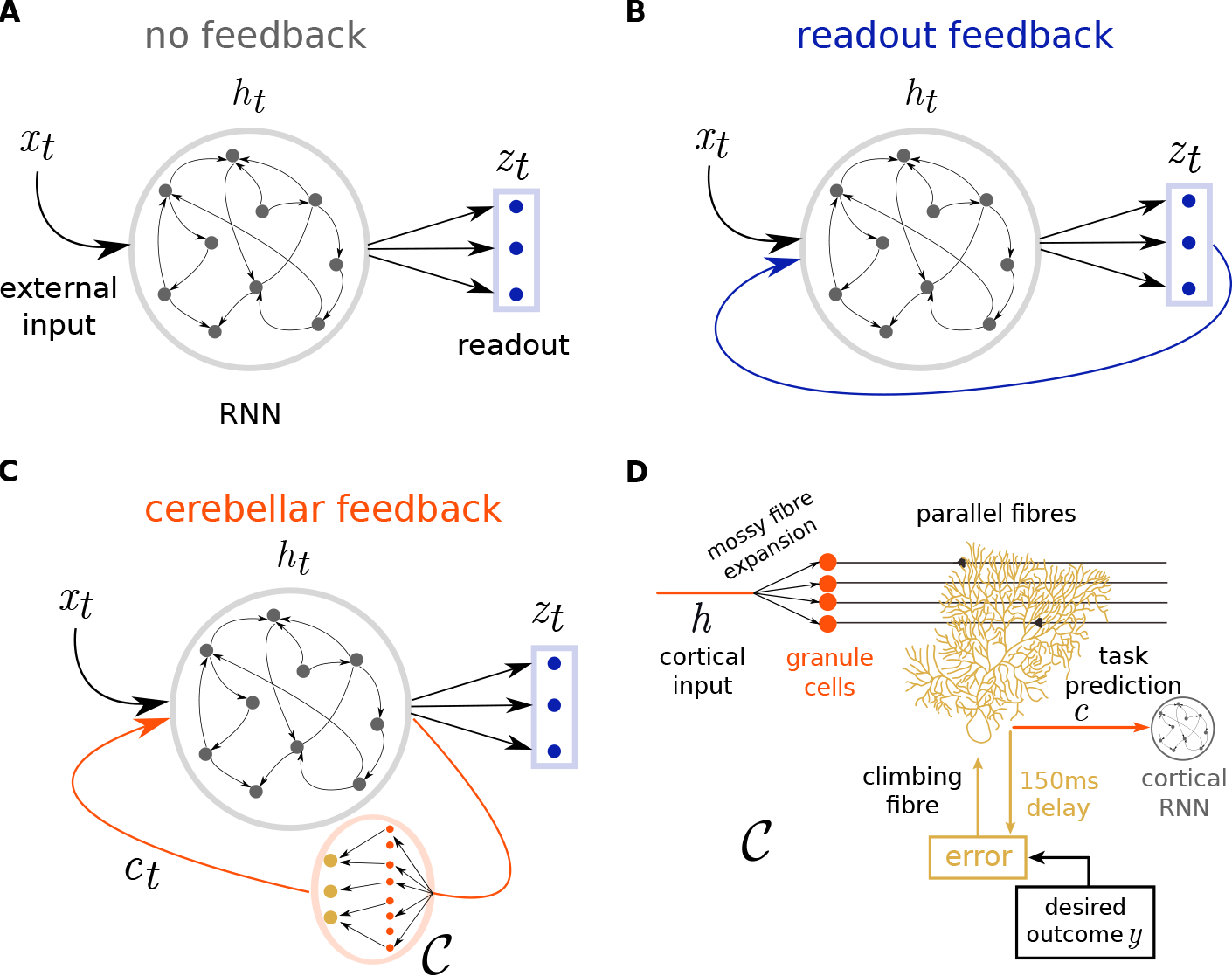
Schematic of cortical recurrent networks with different types of feedback. (**A**) Model variant with *no feedback:* temporal external input (*x*_*t*_) is fed to a cortical RNN (grey) and a linear readout layer (blue) produces the final model output (*z*_*t*_). (**B**) Model variant with *readout-only feedback:* in this scheme there is a feedback loop in which the RNN also receives readout predictions as extra input ^14,16^. (**C**) Model variant with *cerebellar feedback:* a copy of RNN activity (*h*_*t*_) is sent to a (feedforward) cerebellar network 𝒞, which feedbacks to the cortical network its own cerebellar predictions (*c*_*t*_). (**D**) A key property of our cerebellar network is that it learns via behavioural timing-specific learning rules, in line with experimental observations ^32^. In this learning rule the error between the cerebellar prediction *c* and future behavioural outcomes *y* (150 ms) triggers plasticity via climbing fibers at the parallel fibre input of Purkinje cells.

Here we focus on the feedback loop between two key brain regions, the cortex and *the cerebellum*. The cerebellum is a highly plastic system and is well placed to drive cortical dynamics via a set of stereotypical, but functionally separable cortico-cerebellar loops ^20,21^. Indeed, an ever-growing array of clinical ^22^, functional imaging ^23,24^, and optogenetic ^25–27^ studies support an important cerebellar contribution to cortical activity in both motor and non-motor domains. Recently, two hypotheses on the computational role of cortico-cerebellar loops have been put forward ^28–31^. The first asserts that the cerebellum reinforces cortical-dependent goal-directed behaviour by appropriately steering or stabilising cortical states in real-time ^28,29^. The second also promotes the cerebellum as a facilitator of goal-directed cortical transitions, but it does so indirectly via teaching signals which lead to cortical plasticity ^30,31^. Whilst these two views may co-exist, it is the former that is well placed to operate under weakly plastic cortical networks. Moreover, the cerebellum acting as an instantaneous driver of cortical dynamics is in line with the fast activity-dependent cortico-cerebellar interactions that have been observed experimentally ^25–27^.

Here we put forward a computational framework in which the cerebellum learns to rapidly steer and stabilise task-dependent cortical dynamics. We test this model on a variety of motor and non-motor tasks, proposing that the cerebellum is optimised to support task acquisition in the cortex. This reduces the burden of learning in cortical networks and allows a given cortical area to rapidly switch between different tasks. In line with this, we show that a strong cortical dependence on cerebellar feedback arises after learning, consistent with recent behavioural and optogenetic experiments. Finally, we use this model to put forward a cerebellar-to-cortical systems consolidation theory, in which quickly learnt task-specific information encoded by the cerebellum is gradually transferred to the cortex. Overall, we introduce a computationally and experimentally supported theory for cerebellar-supported task acquisition, switching and consolidation in the brain.

## Results

### A computational model of cerebellar-driven cortical dynamics for task acquisition

To study the role that cerebellar feedback can have in driving cortical dynamics during task acquisition, we explore different variants of cortical RNNs: without feedback (Fig. 1A), with readout feedback (Fig. 1B) ^14,16^ and with feedback provided by a cortico-cerebellar loop (Fig. 1C). We introduce a model of cortico-cerebellar loops, in which a *cortical* RNN is reciprocally connected to a feedforward *cerebellar* network 𝒞. In our model, temporal RNN representations **h**_*t*_ are passed onto the cerebellar network to compute task-specific predictions **c**_*t*_, which are then sent back to the same cortical RNN. The final model output **z**_*t*_ is then a linear readout of the RNN activity

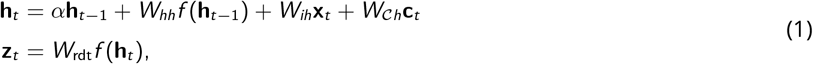

where *α* denotes the cortical internal memory (or leak) of the RNN neurons, *f* (*x*) is the cortical activation function which is set as tanh(*x*). *W*_*hh*_, *W*_*ih*_, *W*_*𝒞h*_ are the recurrent, input, and cerebellar weights onto the RNN respectively, and *W*_rdt_ are the readout weights (see Supplementary Fig. S1 for a detailed schematic). For computational efficiency and due to the relatively long duration of the tasks we train our model using a discrete approximation of a continuous RNN (see Methods). To highlight the need for optimised network connectivity rather than inherent cortical memory mechanisms, in our experiments we generally focus on small *α* = 0.1 (see Methods).

The cerebellar feedback **c**_*t*_ is a feedforward computation *C* on the previous RNN activity

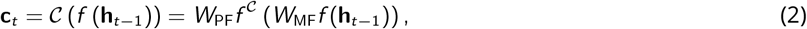

where *W*_MF_ represent the cerebellar (input) mossy fibre (MF) weights onto granule cells (GC) and *W*_PF_ the parallel fibre (PF) weights from GC to Purkinje cells (PC), here representing the output. Together, these constitute the main stages of processing in the cerebellum ^33–35^. In general we model *W*_MF_ as highly divergent with an input/output ratio of 1:20 (see Methods) and *f* ^*𝒞*^ (*x*) as a rectified linear function (ReLU), in line with the large numbers of cerebellar GCs and responses ^36,37^. As we demonstrate in our results, and consistent with prior work, the dimensionality expansion and non-linearity at the GC layer enables better representations during learning.

We use biologically plausible gradient descent ^38^ to optimise cortical weights during the acquisition of a given task (Eq. 1). In particular, we minimise the temporal error *E*_*t*_ = *ℰ*(**z**_*t*_, **y**_*t*_), where **y**_*t*_ denotes the desired task outcome at time *t* and *E* is the task error function (see Methods). These weights can all be optimised simultaneously during learning – we refer to this case as *fully plastic*. However, a key idea that we put forward in this study is that it is not the neocortex, but in fact the cerebellum, which acts as a key driver for task acquisition. For this reason we highlight the case in which RNN plasticity is constrained. In particular, we focus on conditions in which RNN plasticity is either absent – *fixed RNN* case, or in which plasticity is strictly limited to its input synapses (i.e. only *W*_*ih*_, *W*_*𝒞h*_ in Eq. 1 are plastic) – *input plastic* case. The latter case considers both plasticity at sensory and cerebello-cortical input during task acquisition, in line with experimental observations showing plasticity at cerebellar pathways to the cortex ^39,40^.

In contrast to cortical learning, the cerebellum is always optimised, through a separate but related cerebellar error 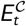. In line with classical models of the cerebellum ^33^ we assume that learning occurs at the parallel fibres *W*_PF_, mediated by climbing fibre error signals, whilst mossy fibres inputs *W*_MF_ remain fixed. Like the cortical prediction error, the cerebellar error function depends on the desired task outcome **y**. However, as we will see later, it is advantageous for the cerebellum to provide predictions of future outcomes. To enable this we formulate a temporal cerebellar learning rule. In this rule the cerebellum learns by comparing its own past output within a predefined time-window *τ*, with current desired outcomes (Fig. 1D), 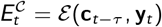 – behavioural timing-specific learning rule. This learning rule then predicts the need for temporally precise coordination between parallel fibre inputs and subsequent climbing fibre error signals to achieve plasticity, in line with experimental findings ^32,41–45^. Therefore, it enables the cerebellum to predict future outcomes effectively, i.e. **c**_*t*_ *≈* **y**_*t*+*τ*_. For our motor-based tasks we generally consider a cerebellar time window of *τ ≈* 150ms ^32^ and for the later cognitive tasks use longer windows *τ ≈* 600ms (see Methods).

### Cerebellum learns to drive cortical dynamics during a line drawing task

To study the functional consequences of cortico-cerebellar loops we first test the model in a motor-based line drawing task. In this task the model receives one out of six cues at the beginning of the task and learns to either remain still or produce one out of five possible straight lines (Fig. 2A; see Methods). Feedback provided by desired outcomes (i.e. straight lines) is provided at each timestep. Consistent with behavioural studies on cerebellar patients ^46^, we find that cerebellar feedback significantly improves learning of the task and final performance (Fig. 2A,B). The ability for cerebellar feedback to facilitate learning does not depend on the degree of plasticity and internal memory in the cortical RNN (Fig. 2C). Interestingly, a fixed RNN with a plastic cerebellum achieves the same learning performance as a fully plastic or input plastic RNN. In contrast, when no feedback or a simple readout feedback is provided the network can fail to learn the task due to the leaky properties of RNNs (Fig. 2B,C). Classical cerebellar models pose that the cerebellum can act as a direct controller of motor tasks ^33^. To contrast this view with our model we also train an RNN with a direct cerebellar readout, which apart from the cortico-cerebellar feedback weights uses the same free network parameters, and find it insufficient to learn the task (Supplementary Figs. S1 and S2).

**Figure 2.**
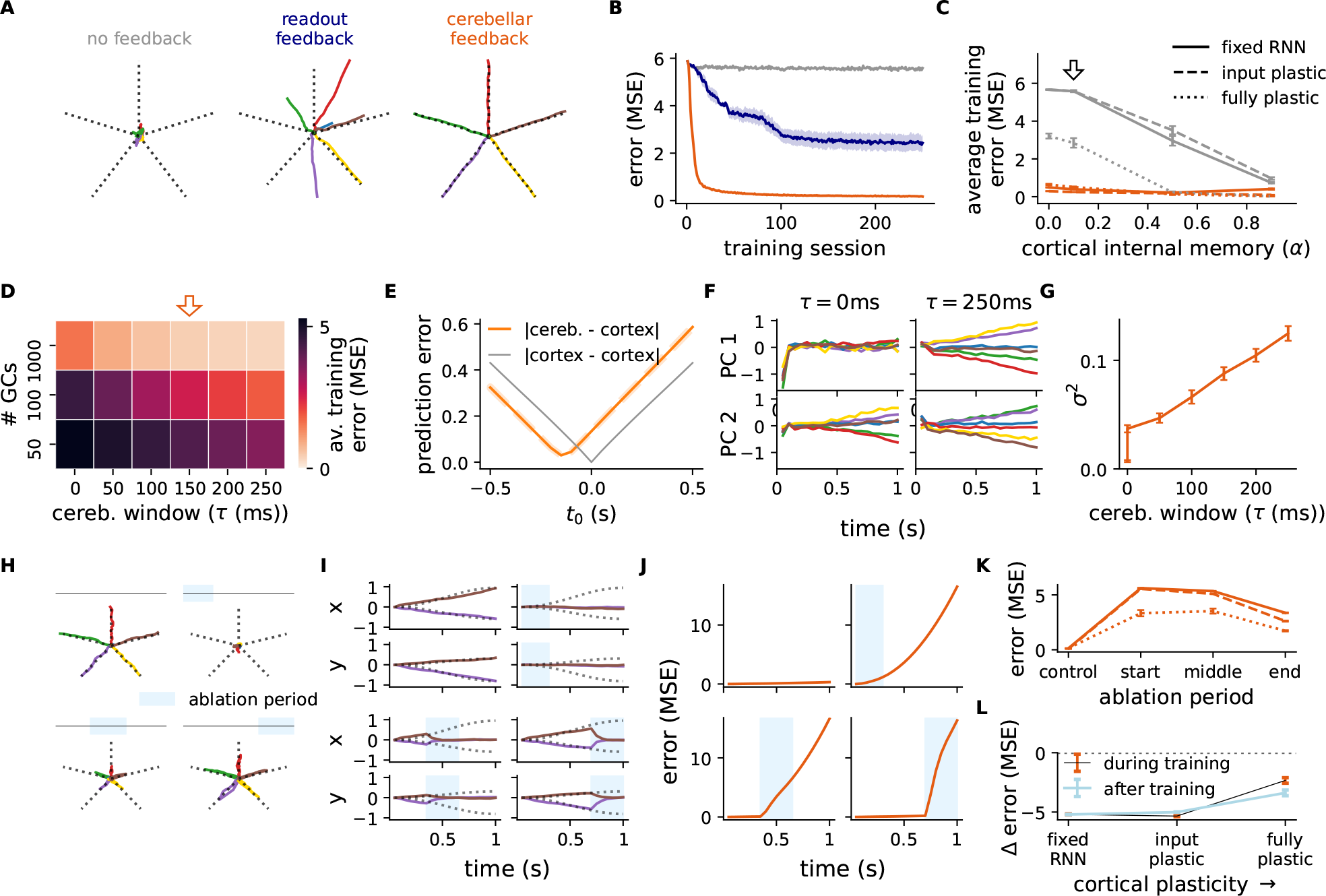
Cerebellum learns to drive cortical dynamics during a line drawing task. (**A**) Given one of six possible stimuli at the first timestep the model must learn to draw a corresponding line (dotted black line) or remain still. Model output after training is shown for three model architectures with a fixed RNN. (**B**) Learning curves of models in A (same colour-coding). MSE denotes mean squared error. (**C**) Average training error across different levels of RNN internal memory (*α*) and plasticity (fixed RNN, input plastic and fully plastic) for the no feedback and cerebellar feedback models; arrow denotes cortical internal memory used in the other panels (*α* = 0.1). (**D**) Average training error of cortico-cerebellar model under varying numbers of granule cells and cerebellar temporal windows (*τ*). Orange arrow denotes default parameter choices. (**E**) Prediction error between cortical output and itself (gray) or cortical output and cerebellar output (orange) for different temporal delays. (**F**) Evolution of first (upper panel) and second (lower panel) principal components of cortical RNN for different stimuli, colour-coded as in (A) using small (*τ* = 0ms) and large (*τ* = 250ms) cerebellar time windows. (**G**) Variance across cues from both first and second PCs (cf. F) for different cerebellar temporal windows, *τ*. (**H**) Model output for different periods of cerebellar ablation (blue box represents period of ablation). (**I**) Output *x* and *y* coordinates of the lines drawn in H. (**J**) Average model error across all inputs for ablation periods in H,I. (**K**) Average error for different degrees of plasticity and ablation periods (left to right) as in H-J. (**L**) Average change in task error for models with versus without cerebellar feedback during (black) and after (blue) training for different degrees of cortical plasticity. All results are averaged over 5 different initial conditions. Error bars represent standard error of the mean.

Next, we study how two known cerebellar features: (i) a large number of granule cells and (ii) behavioural timing-specific plasticity rules contribute to task proficiency. We find that a combination of high numbers of granule cells with a learning rule with a non-zero temporal horizon, *τ*, result in better cerebellar learning (Supplementary Fig. S3), which in turn drives better cortical representations and overall task performance (Fig. 2D and Supplementary Figs. S3,S4). Moreover, because both the cortical RNN readout and cerebellar network are trained on the same desired outcome, we observe that cerebellar output effectively predicts cortical readout *τ* ms ahead (Fig. 2E). Our model thus provides a theory of how the cerebellum learns to predict upcoming movements ^47,48^.

The advantage of a large number of granule cells has been well studied is likely due to better linear separability of its inputs ^49^. However, what are the computational advantages of the cerebellum providing the cortical RNN with expected future outcomes? Due to RNN leakiness, sensory cues are rapidly forgotten. Therefore a high cerebellar *τ* gives the cerebellar network the ability to map RNN activity to desired outcomes early on in the task. Consistent with this we find that the predictive cerebellar output drives outcome-dependent RNN representations (Fig. 2F,G). This result showing potent initial drive of cortical activity could provide a justification for the observed role of the cerebellum in movement initiation ^50,51^.

Finally, to directly examine the role of cerebellar feedback on cortical dynamics, we inhibit - or “ablate” - cerebellar output (i.e. **c**_*t*_ = 0 in Eq. 1) during different stages of the task. In each case we observe significant impairment in the model output which returns to baseline (Fig. 2H-J). Moreover, this effect is most detrimental to task performance when ablation occurs at the start (Fig. 2K). These findings are consistent with the observed freezing effect of cerebellar lesions on gait ^52^. In line with both cortical and cerebellar networks working jointly to perform the task, we find that when the RNN is fully plastic cerebellar ablations have a significant but reduced impact on the cortical dynamics (Fig. 2K,L and Supplementary Fig. S5). We also observe that the cortical RNN is particularly sensitive to the presence of noise in cerebellar output. When noise is added to its output it leads to irregular behaviour (Supplementary Fig. S6), in line with the classical motor symptoms of cerebellar ataxia ^53^.

Taken together, this motor-based task highlights the computational benefits of training a cerebellar network to drive cortical dynamics, predicting that the cortex can critically depend on cerebellar feedback for successful task execution. Furthermore, we demonstrate that cerebellar plasticity can effectively replace the need for local cortical plasticity.

### Cerebellar-mediated task switching in cortical networks

We have shown that cortico-cerebellar loops can enable successful task learning with minimal cortical plasticity. This opens the possibility of reusing cortical networks across different contexts and behaviours.

To demonstrate the model’s ability to adapt and perform context-dependent task switching, we consider how models trained in the line-drawing task can be retrained to a curl-field variant ^54^. In particular, we analyse how the cerebellar network can (i) successfully enable learning in a new task context and also (ii) rapidly revert, or *switch*, to a previously learned context.

As expected, when the new task context is introduced to the model, there is a steep increase in error before the model successfully learns the new task (Fig. 3A, left and middle). Notably, however, when the original task is reintroduced, the fixed RNN model recovers the initial dynamics significantly faster than the fully plastic model and more faithfully captures the behavioural data from macaque monkeys ^54^ (Fig. 3A, right). This relatively slow switching back suggests that the fully plastic RNN is more prone to forgetting the original task ^9^.

**Figure 3.**
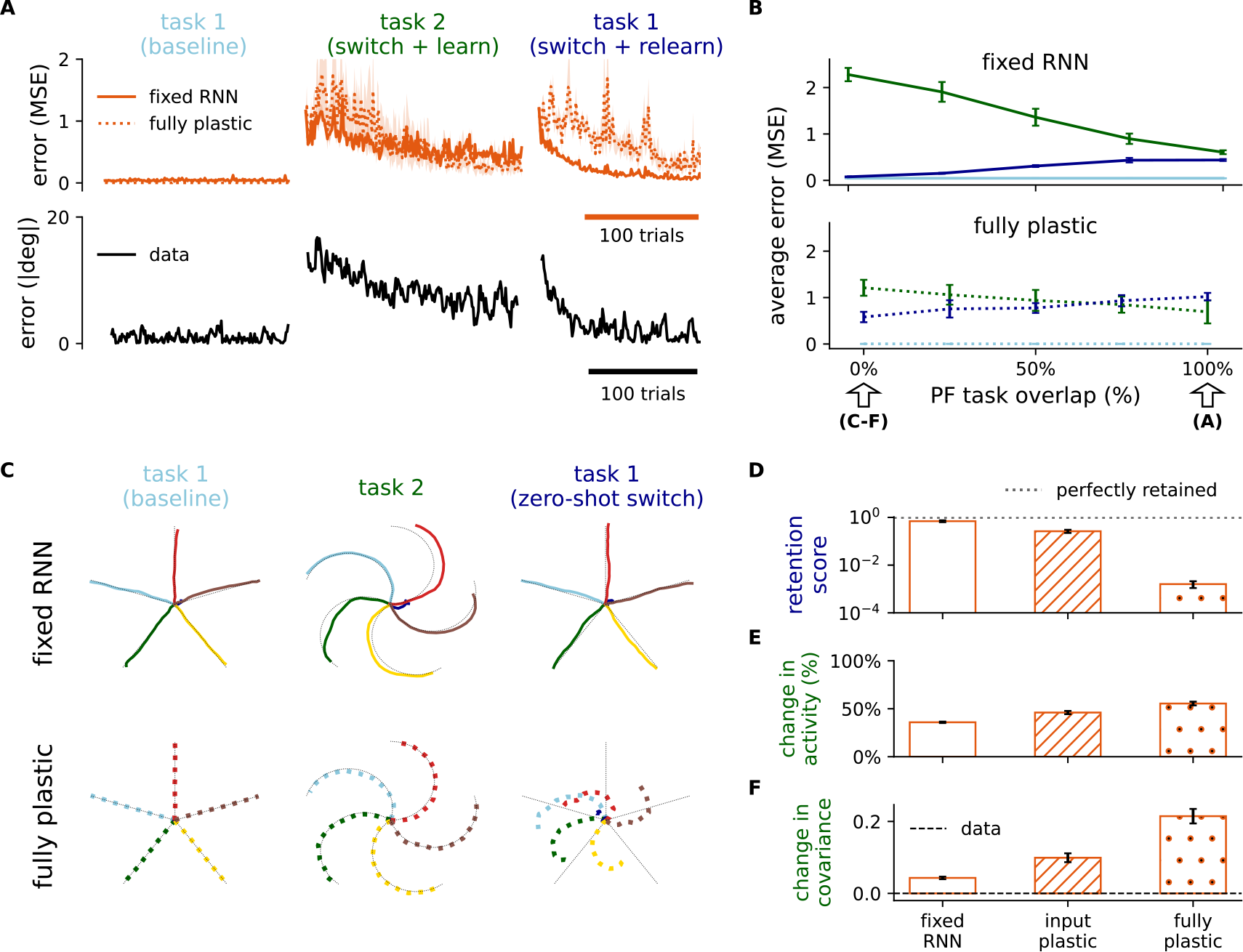
Context-dependent cerebellar feedback can enable multi-task learning and switching in the cortex. (**A**) Training error of cortico-cerebellar models originally trained for line drawing (cf. Fig. 2; *α* = 0.5). The models continue to execute the line-drawing task (left) before being trained on a novel curl-field variant of the task (middle) and then finally switch back to the original task (right). Data from behavioural experiments in macaque monkeys is reproduced here for comparison (bottom; ^54^). (**B**) Average training error across different levels of parallel fibre (PF) task overlap for the different tasks for the fixed RNN (top) and fully plastic (bottom) models. Task periods colour-coded as in A. Arrows denote degree of PF task overlap used in A and C-F. (**C**) Model output for each of the three training periods defined in A for the zero-overlap condition; “zero-shot” output corresponds to the model output in the first trial when task 1 is reintroduced. (**D**) Model retention score for task 1. The retention score is computed as the error of task 1 during baseline over the error at the first trial after switching back to task 1. (**E, F**) Change in (E) activity and (F) covariance in the RNN population between task 1 (baseline) and after learning task 2. Mean changes in experimental data in F are reproduced (see Methods) from neuronal recordings obtained from premotor (PMd) and primary motor (M1) cortices in macaque monkeys ^54^. All results are averaged over 5 different initial conditions. Error bars represent standard error of the mean.

We then asked how the cerebellar network might enable even faster task switching. In line with observed context-dependent activations ^55,56^ and plasticity rules ^57^ in the cerebellum, we consider cerebellar PFs which are task-specific. The extent of task-specificity at PFs is modelled by the *PF task overlap*; full overlap (100%) would imply that the same exact PFs are used across task contexts, while zero overlap (0%) implies that a completely different set of PFs is used for each task respectively.

Our results show that the degree of PF task overlap predicts a tradeoff between the speed of learning the new task and the ability to rapidly switch back to the original task (Fig. 3B). Specifically, whilst maximal PF task overlap is beneficial when a new task is introduced, rapid switching is favoured when distinct PFs are used. To highlight the ability to immediately switch back to the original task (zero-shot switch) we focus on the zero-overlap case. For the fixed RNN, but not the fully plastic RNN, the model achieves near-perfect switching to the original task (Fig. 3C,D). Consistent with the need to learn a new task all models show a substantial change in the neuronal activity (Fig. 3E and Supplementary Fig. S7A). However, we expect that models with minimal local cortical plasticity should result inminimal changes in the underlying dynamics of both tasks. To test this, we measure changes in the the covariance of the neuronal activity between the new task and the initial task (see Methods and ^58^). As predicted, only the models with reduced cortical plasticity show the minimal changes observed experimentally (Fig. 3F and Supplementary Fig. S7B). On the other hand, for the fully plastic model the dynamics acquired after switching back to the initial task are significantly different to baseline (Supplementary Fig. S7C,D). This suggests that the fully plastic model learns a new solution to the initial task, explaining its relative slowness in switching.

Overall, we apply our models to demonstrate a cerebellar-driven solution to multi-task learning and task switching. We show that the underlying dynamics preserved by a fixed cortical RNN, supported by context-dependent cerebellar feedback, can support rapid behavioural changes whilst minimising forgetting of previously acquired task knowledge.

### Cerebellar temporal basis supports non-linear drawing task

Above we have modelled a case in which the cerebellum learns to drive cortical dynamics using a specific predictive time-window (namely *τ* = 150ms). However, a recent study has revealed a diversity of temporal plasticity windows to be at play in the cerebellum ^32,59^ (Fig. 4A). Such diversity of temporal windows may enable the cerebellum to learn a *temporal basis* for upcoming events, which may enhance the cerebellum’s ability to predict future outcomes.

**Figure 4.**
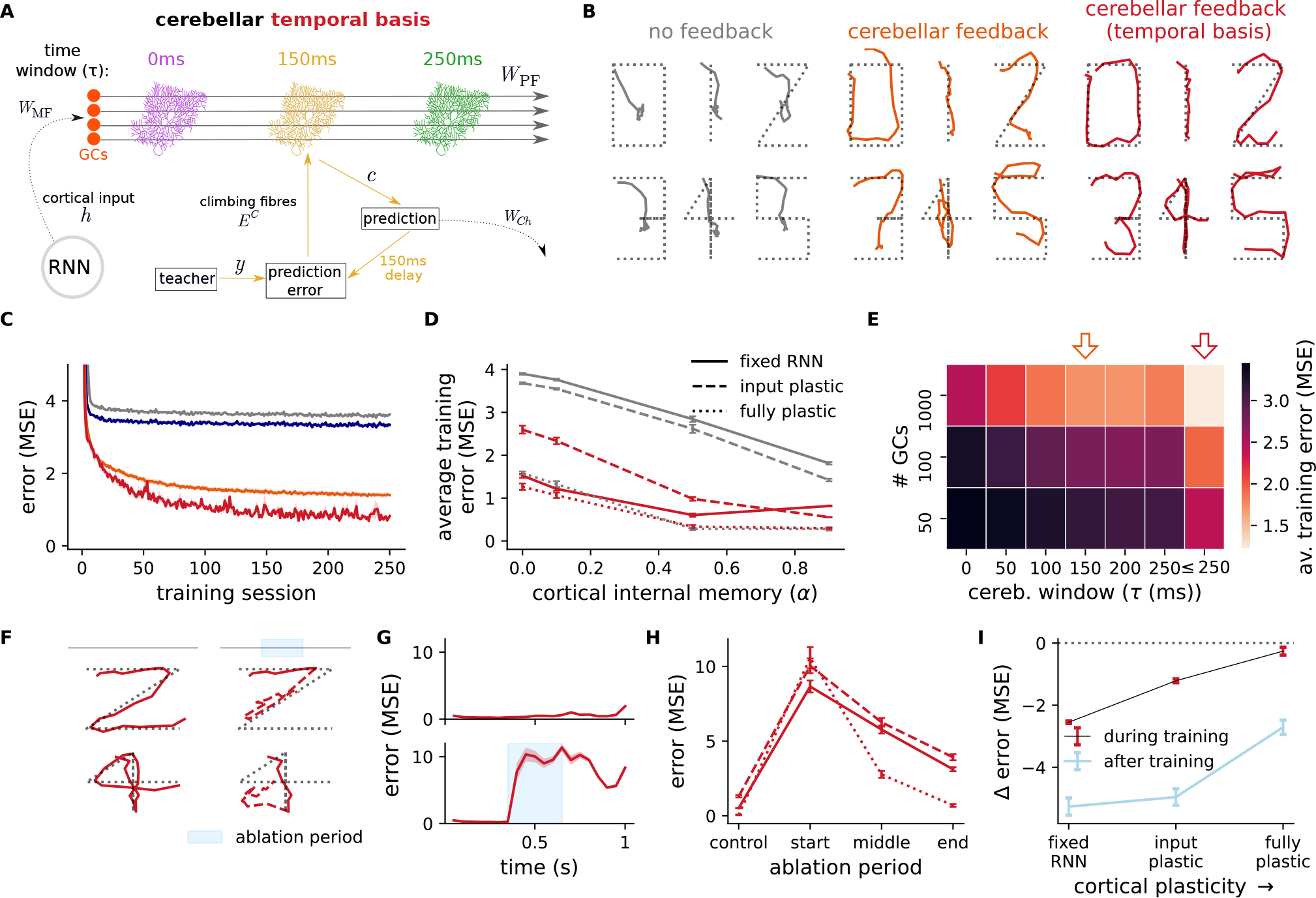
Cerebellar temporal basis supports cortical dynamics of a non-linear digit drawing task. (**A**) Schematic of cerebellar learning with a temporal basis. We consider multiple populations of Purkinje cells with different learning time windows *τ*. (**B**) Model output after training for different input examples of the digit drawing task (fixed RNN; *α* = 0.1). (**C**) Learning curves of models in B together with readout feedback model (blue). (**D**) Average training error across different levels of RNN cortical internal memory (*α*) and plasticity assumptions. (**E**) Performance of cerebellar feedback for different numbers of granule cells and and cerebellar time windows. Orange arrow indicates default parameter choices with a single cerebellar time window; red arrow indicates temporal basis model with multiple time-windows. (**F**) Model output under control and cerebellar ablation conditions for example inputs (digit 2 in upper panels and digit 4 in lower panels); dashed red line represents model output during and after ablation period. (**G**) Average model error across all inputs for control (left) and ablation (right) conditions. (**H**) Average error for different degrees of cortical plasticity and ablation periods (middle period illustrated in F,G). **I**, Average change in task error for models with versus without cerebellar feedback during (black) and after (blue) training across different degrees of cortical plasticity. All results are averaged over 5 different initial conditions. Error bars represent standard error of the mean.

To demonstrate the benefit of diversity in temporal windows we consider a more realistic (and challenging) variant of the line-drawing task in which the model is now trained to produce a digit-like output (Fig. 4B; see Methods). This task is selected so as to produce a non-linear and highly varied set of future desired outcomes and therefore the need for richer cerebellar predictions. In particular, we consider a cerebellar network which simultaneously learns with a range, or “temporal basis”, of time-windows *τ*_*i*_ ∈ [0ms, 250ms] such that its prediction effectively spans a relatively long window of upcoming desired outcomes (see Methods).

We find this heterogeneity of cerebellar time windows to enable both faster learning and higher performance thresholds (Fig. 4B,C and Supplementary Fig. S8). As expected, when considering the simpler line-drawing task having multiple time windows does not improve learning (Fig. S8C). Moreover, in line with the results above, a fixed RNN achieves a performance comparable to the plastic RNN models across different degrees of internal memory in the cortical network (Fig. 4D). When comparing the network performance across different numbers of granule cells and time-windows, we find that higher numbers of granule cells combined with multiple time-window learning achieves the best average learning performance (Fig. 4E). Finally, as with the simpler line-drawing task, we find that cerebellar ablation is detrimental to the maintenance and development of these representations (Fig. 4F-H) in a way that depends on the degree of cortical plasticity (Fig. 4I and Supplementary Fig. S9).

These results suggest that the diversity of behavioural-specific learning windows observed experimentally in the cerebellum ^32,59^ improve behaviour when in the presence of more challenging task conditions.

### Cerebellar-driven cortical dynamics maintains beliefs in an evidence accumulation task

So far we have focused purely on motor-based tasks, but growing evidence strongly suggests that the cerebellum also plays important roles in functions that go beyond direct motor control ^21,60,61^. To demonstrate this we model an evidence accumulation task that has been shown to be cerebellar-dependent ^26^. In this study Deverett et al. ^26^ showed that optogenetic inhibition of the cerebellar output nuclei disrupts the ability of mice to determine whether the left or right cheek received more air puffs over a period of time (Fig. 5A). Unlike the previous tasks, here the desired outcome is only provided at the end of the task, making error-related signals highly sparse.

**Figure 5.**
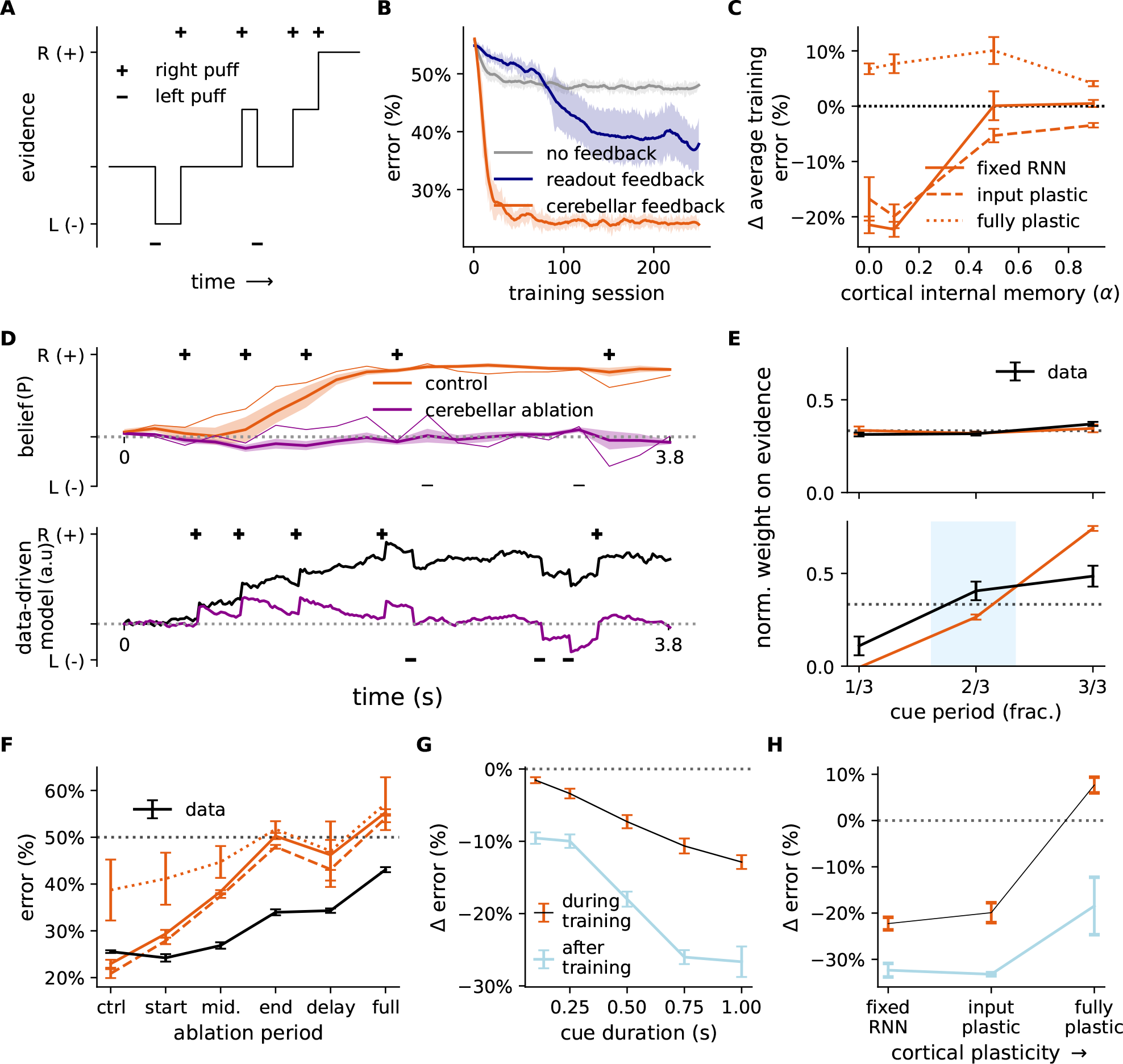
Cortico-cerebellar model mimics mouse behaviour during evidence accumulation task. (**A**) Schematic of evidence accumulation task ^26^: a random sequence of non-zero inputs (“air puffs”) is delivered in the leftward (-) or rightward (+) direction. The model must integrate this input and decide at the end of the task which side received more input overall. (**B**) Learning curves of models (fixed RNN; *α* = 0.1) without feedback (grey), with readout feedback (blue) and with cerebellar feedback (orange). (**C**) Change in average training error of the cortico-cerebellar model with respect to the no feedback model across different levels of cortical internal memory (*α*) and degrees of cortical plasticity. (**D**) Model beliefs over time without (orange) and with complete cerebellar ablation (purple) in model (upper panels) and data-derived behavioural model (lower panels) reproduced from Deverett et al. ^26^. Thin model lines represent one example seed. Belief *P* denotes model output probability. (**E**) Normalised regression weights at different periods of input presentation (cue) during control (upper) and ablation (lower) conditions for both model (orange line) and behavioural data (black line). (**F**) Model and data error under different ablation periods and degrees of cortical plasticity. (**G**) Average change in task error for models with versus without cerebellar feedback across different cue durations. (**H**) Average change in task error for models with versus without cerebellar feedback during and after training across different degrees of cortical plasticity. All model results are averaged over 5 different initial conditions. Error bars represent standard error of the mean.

Similar to the motor tasks studied above, cerebellar feedback improves task learning relative to models without feedback or with readout feedback (Fig. 5B). Moreover, a fixed RNN achieves performance comparable or even superior to the fully plastic models across a range of degrees of cortical internal memory (Fig. 5C and Supplementary Fig. S10). These results suggest that weakly plastic cortical networks driven by the cerebellum may also be sufficient for learning cognitive-based tasks with sparse error information.

Next, our ablation analysis reveals strong similarities to the optogenetic observations by Deverett et al. ^26^. In particular, cerebellar ablation greatly impairs the model’s capacity to maintain and develop beliefs, mirroring the behavioural effects observed experimentally (Fig. 5D and Supplementary Fig. S11). Indeed, using the same behavioural regression performed by Deverett et al. ^26^ (see Methods), we show that cerebellar ablation in latter periods leads to a final choice in which information about previously seen inputs is greatly reduced (Fig. 5E), in line with experimental findings. Because more information is effectively lost, we find that ablation near the end of the task has a particularly detrimental impact on task performance, consistent with behavioural observations (Fig. 5F), and this leads to a sub-chance ability to perform “history-centric” trials which rely more on initial inputs (Supplementary Fig. S11; see Methods). These ablation results also emphasize that even though the cerebellum is trained with teaching signals close to the end of the task, cerebellar predictions prove to be valuable earlier in the task (Fig. 5F). Finally, to demonstrate that task performance also depends on cortical dynamics, we performed (partial) ablation to cortical RNN and observed similar behavioural deficits (Fig. S12A-C).

Given that cerebellar feedback is necessary to preserve information over time and avoid leaky cortical dynamics, we predicted that the behavioural effect of cerebellar ablation would depend on the timescale of the task and would weaken for shorter task durations. Indeed, we find that the performance effect of ablation increases as a function of task length (Fig. 5G and Supplementary Fig. S13; see Methods). Like in the previous motor-based tasks, our model predicts that cerebellar feedback is particularly helpful when in the presence of weak cortical plasticity (Fig. 5G,H).

Overall, our model predicts that the proper maintenance of model selectivity depends critically on cerebellar feedback during evidence accumulation. Consistent with behavioural results, these effects are emphasised when cerebellar ablation occurs in the later stages of the task.

### Cerebellar feedback sustains cortical dynamics in a delayed association task

Next we aim to demonstrate that cerebellar networks can also effectively drive cortical dynamics in tasks with long delay periods, while capturing both neuronal and behavioural observations. To achieve this we model a delayed association task which was recently shown to dependent on cortico-cerebellar loops ^25^. In this study mice were presented with one of two stimuli (left or right) followed by a delay period, after which they were trained to lick in the corresponding direction (Fig. 6A, top). At the same time neural selectivity was recorded both in the anterior lateral motor cortex (ALM) - a working memory and planning region - as well as the cerebellar output nuclei (Fig. 6A, bottom). Timed photoinhibition was used to reveal ALM selectivity to strongly depend on the cerebellar output nuclei, and vice versa.

**Figure 6.**
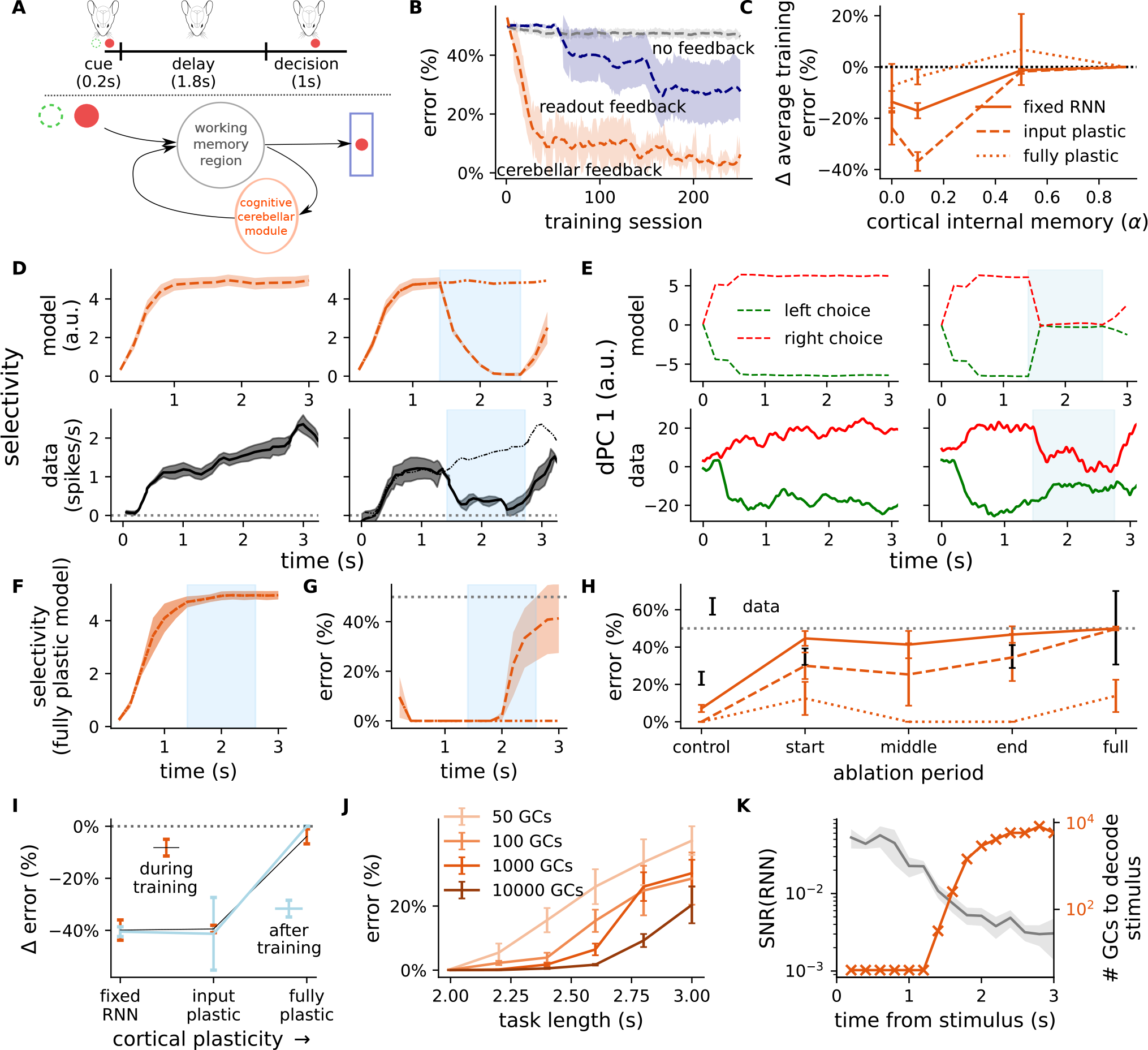
Cerebellar network sustains cortical dynamics during delayed association task in line with optogenetic experiments. (**A**) (Top) Delayed association task; a sensory cue is presented followed by a delay and decision period ^25^. (Bottom) The cortico-cerebellar loop models the interactions between a working memory region and a cognitive module of the cerebellum. (**B**) Learning curves of model without feedback (grey), readout feedback (blue) or cerebellar feedback (orange) for models with an input plastic RNN (*α* = 0.1). (**C**) Change in average training error of the cortico-cerebellar model with respect to the no feedback model across different levels of cortical internal memory (*α*) and degrees of plasticity in the cortical RNN. (**D**) Cue selectivity during the delay period without (left) and with cerebellar ablation (right; blue area denotes period of ablation and thin line shows control) in the model (upper panels) and optogenetic experiments (lower panels) reproduced from Gao et al. ^25^. (**E**) First decision principal component (dPC) during the delay period without (left) and with (right) cerebellar ablation in the model (top) and in optogenetic experiments (bottom) ^25^. (**F**) Cue selectivity during the delay period with cerebellar ablation when using the fully plastic RNN (cf. D). (**G**) Model error during cerebellar ablation (input plastic RNN; control error shown with dashed-dotted line). Dotted grey line denotes chance level. (**H**) Average error from cerebellar ablation at different points during the delay period and different degrees of cortical plasticity. (**I**) Average change in task error for models with versus without cerebellar feedback during and after training across different degrees of cortical plasticity. (**J**) Model error for different numbers of cerebellar granule cells (GCs) and delay period lengths in the delayed association task (fixed RNN; *α* = 0.1). (**K**) Signal-to-noise ratio (SNR) of RNN activities (left y-axis) and number of GCs needed to decode the stimulus from these activities (right y-axis). Results are averaged over 5 different initial conditions. Error bars represent standard error of the mean.

To model this task we follow the same protocol used experimentally ^25^, where one of two possible cues are presented followed by a delay period, after which the model makes a cue-based response (left or right; see Methods). Given the lack of sensory or teaching information during the delay period the cortico-cerebellar network it is particularly vital in this task to sustain stimulus representations. It is important to note that a standard randomly initialised RNN is unlikely to achieve this property, since memories of previous inputs naturally decay in the absence of task-induced plasticity ^19^.

We observe that cerebellar feedback consistently enables task acquisition (Supplementary Fig. S14), and identify a particularly interesting case when plasticity in the RNN is limited strictly to its input synapses (input plastic). In this case cerebellar feedback significantly improves cortical learning to reach near-perfect performance, whilst also enabling a high degree of stability in task selectivity throughout the delay period (Fig. 6B-D and Supplementary Fig. S14). We speculated that for this task input plasticity is particularly important, because the cerebellum is required to sustain task-specific predictions in the RNN throughout the entire delay period. We verified this stronger cerebello-cortical drive by using concepts from control theory ^62^. In particular, we can explicitly relate cerebello-cortical optimisation to a quantitative increase in the impact, or *energy*, of cerebellar feedback onto RNN activity (Supplementary Fig. S15; see Methods). Moreover, the ability of the cerebellum to drive cortical dynamics should depend on the cortical network’s ability to express those dynamics. In line with this view our results show that (even untrained) cortical recurrent weights are important in maintaining cerebellar predictions over time (Supplementary Fig. S16).

Next, to demonstrate that the cerebellum helps drive task-specific dynamics in the cortical RNN we performed a simulated ablation in which the cerebellum is transiently removed during the delay period. Consistent with *in vivo* neural recordings ^25^, we find that both cerebellar and cortical ablation drastically disrupts cortical task selectivity (Fig. 6D and Supplementary Fig. S12D-F). We next show a similar effect in the model’s latent dynamics: using demixed principal component analysis ^63^ we observe that the choice component of the RNN’s population dynamics collapses rapidly during the ablation period, consistent with neural data (Fig. 6E). As with the previous tasks, our model predicts that this effect depends on the degree of plasticity in the cortical RNN. In particular, a fully plastic RNN notably fails to capture the strong dependence on cerebellar feedback as observed experimentally (Fig. 6F and Supplementary Fig. S17; compare with Fig. 6D, bottom right). Indeed, we only observe an effect on performance consistent with experimental findings when cortical plasticity is limited (Figs. 6F-I). Taken together our results suggest that the cerebellum, not the cortex, is the primary site of learning during the acquisition of this working memory task ^25^.

Overall, these results demonstrate that our model can capture working memory tasks and the observed dependency of cortical dynamics on cerebellar input. Moreover, our model makes the prediction that the cerebellum is a key site of plasticity during acquisition of delayed association tasks.

### Cerebellar divergence decodes task-relevant signals from cortical memory

As mentioned, a prevalent feature in classical cerebellar theories is that the divergence provided by the granular layer enables a linear separation of similar inputs ^34,35,64^. Whilst this has typically been studied using isolated models of the cerebellum, it has recently been suggested that this feature may be of relevance in the context of memories in the cortex which merge or “collapse” onto similar representations over time ^65^.

We tested this in our model and observed that a large quantity of cerebellar granule cells is indeed particularly valuable when the initial stimulus is followed by a long delay (Fig. 6J). In particular, our results show that as the signal-to-noise ratio (SNR) of the cortical RNN activity decreases over time, more granule cells are required to decode the stimulus from that activity (Fig. 6K; see Methods). The model therefore demonstrates that the cerebellum is uniquely placed to decode cortical representations whose task-relevant signals naturally weaken over time. This may explain recent experimental results which suggest the cerebellum is particularly important for tasks which induce long delay periods ^66^.

### Cerebellar task knowledge can be consolidated in the cortex

In each of the previous tasks, cerebellar feedback is shown to mediate learning and the maintenance of task-specific cortical dynamics. However, the neocortex is known to encode long-term representations of tasks ^10^. This suggests a need for a “consolidation” period, during which the memory stored in the cerebellum may be transferred to cortical areas.

To demonstrate cerebellar-to-cortical systems consolidation in our model we develop consolidation-specific learning rules. To achieve consolidation we train cortical recurrent weights to mimic cerebellar input (see Methods). In principle, this should be readily attainable, since the addition of cortico-cerebellar feedback itself can be interpreted as a low-rank modification of the RNN weights ^67^. We also gradually decay the cerebellar-to-cortical input weights so that over training the cerebellum stops driving the cortical network, thereby giving full control of the task to the cortical RNN (Fig. 7A).

**Figure 7.**
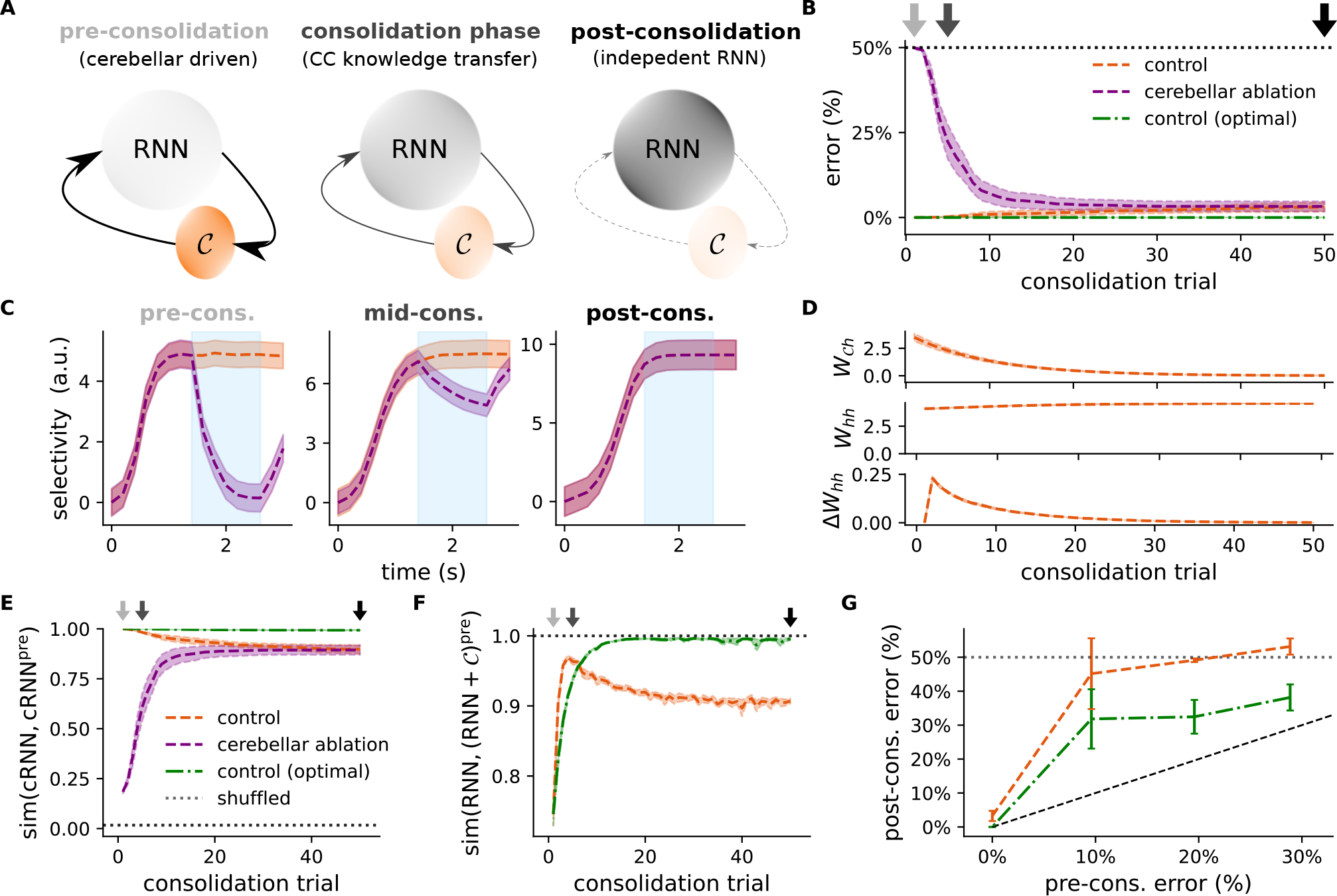
Cerebellum can mediate task consolidation in the cortex. (**A**) Schematic of proposed theory of cerebellar-to-cortical task consolidation. During the initial learning phase (left), task representations are primarily driven by the cerebellum and RNN connectivity is not yet task-specialised. During the consolidation phase there is a period of cerebellar-to-cortical (CC) task information transfer (middle), whereby CC interaction drives plasticity in the cortical RNN. After consolidation (right), the RNN can operate effectively without the need for cerebellar input. The colour of the structures reflects the importance of each component throughout consolidation. (**B**) Model error in the delayed association task (Fig. 6) throughout consolidation with (purple) and without (orange) cerebellar ablation. For reference an optimal consolidation model is also given (green). Dotted black line denotes chance. (**C**) Model selectivity with and without cerebellar ablation at different stages of the consolidation process; titles colour coded according to arrows in B. (**D**) Strength of the cerebellar-to-cortical weights (*W*_*𝒞h*_; top), local cortical weights (*W*_*hh*_; middle) and change in local cortical weights (Δ*W*_*hh*_; bottom) over the period of consolidation. Strength and change is measured by the Euclidean norm. (**E**) Cosine similarity between cRNN (RNN and cerebellar network) activities before and during consolidation. (**F**) Cosine similarity between the learned recurrent input currents (generated locally in the cortical RNN) during consolidation and the total cortical input current (generated locally and by cerebellar-cortical input) in the pre-consolidation network. Similarity of the consolidation model is shown in orange and the optimal consolidation model in green. (**G**) Task error after the consolidation period for models with different initial degrees of performance prior to consolidation. Results are averaged over 5 different initial conditions. Error bars represent standard error of the mean.

We tested this computational theory of consolidation on the cortico-cerebellar models (input plastic condition) trained on the previous delayed association task (Fig. 6D, top left). We consider two types of learning rule. The first is a simple biologically plausible rule, which depends on the ratio of cerebellar-to-cortical input and total RNN activity. Specifically, the recurrent weight *w*_*ij*_ from cortical neuron *i* to *j* evolves according to 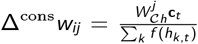 where 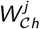 is the *j*th row of *W*_*𝒞h*_ and the denominator a normalising factor, which may be computed by cortical interneurons ^68^. For comparison we also consider a theoretically optimal (but biologically unrealistic) rule based on a least squares solution (see Methods).

In both cases, we observe that the RNNs gradually learn to perform the task without the need for cerebellar input (Fig. 7B,C). During this period, the cerebello-cortical weights decay gradually to zero, whilst relatively small but important weight modifications take place within the cortical RNN (Fig. 7D). By construction of the learning rule, the cerebello-cortical activities throughout the consolidation period closely resemble, or “replay”, their original pre-consolidation values, and the RNN is eventually able to independently recreate the (pre-consolidation) cerebellar-dependent dynamics (Fig. 7E,F). Such “replay” of task-dependent dynamics is consistent with experimental observations of cerebello-cortical interactions during sleep ^69^.

We also find that a model with fixed RNN connectivity does not perform as well as the input plastic condition (Supplementary Fig. S18). This is likely due to better network stability when in the presence of the input plastic, compared to the purely fixed RNN (Figs. S17A,B and Supplementary Fig. S19). Related to this, we find that models which have not yet perfected the task exhibit worse performance after consolidation (Fig. 7G).

In summary, the framework we introduce here suggest that the cortico-cerebellar loops may play an important role in systems consolidation by gradually transferring the rapidly learnt cerebellar knowledge to the cortex.

## Discussion

Growing experimental evidence suggests that cortico-cerebellar loops support behaviour, but their computational roles have remained unclear. Here we have introduced a systems-level modelling framework in which a feedforward cerebellar network receives the state of a cortical RNN and provides task-specific predictions in return. In our model, cerebellar feedback facilitates learning by shaping the underlying cortical dynamics during motor and cognitive tasks in a way that is consistent with both behavioural and optogenetic studies. Our work suggests that the cerebellum is a key site of learning in the brain, allowing for rapid context-switching of cortical dynamics that underlie behaviour. We finish by introducing a theory of cerebellar-to-cortical system consolidation, in which task-specific knowledge is gradually transferred to the cortical network.

Our model is related to previous network architectures in that it uses feedback to enhance neuronal representations and selectivity in an otherwise fixed RNN, thereby facilitating task-relevant downstream processes ^15,16^. There is a growing interest in neuroscience on the role that feedback can play in cortical circuits. For example, two recent theoretical studies demonstrate how thalamic feedback implemented by cortico-thalamic loops can flexibly prepare and execute motor sequences ^17,18^. We highlight two key computational differences in our work. First, in our model feedback is not derived by a linear function of the RNN (as usually done when using simple readout or thalamic networks), but from a divergent cerebellar-like feedforward network (Fig. 1). Second, our model incorporates behavioural timing-specific learning rules in line with experimental findings ^32,41,43–45^. We show that these cerebellar features improve task-acquisition against a standard readout feedback architecture ^14–16^ (Figs. 2, 4, 5 and 6).

By retraining cortico-cerebellar networks in a novel task we propose a key role of the cerebellum in task switching (Fig. 3). In particular, we show that cerebellar feedback may provide a solution to the problem of context-dependent adaptation, which requires (i) an ability to learn a new context but also (ii) an instant retrieval of appropriate response to previously learned contexts ^70,71^. Interestingly, we observe that while recurrent cortical plasticity enables adaptation to a new task context there is catastrophic forgetting of the original context. This is at odds with well-known behaviour in the primate, and provides a computational explanation for why local modifications in the monkey cortex during motor adaptation appear to be limited ^54^. In our model rapid task switching is achieved by context-specific activation of cerebellar parallel fibres. In future work it would be of interest to compare different mechanisms by which the cerebellum may realise context-dependent processing; for example, a recent study has suggested that dendritic gating via cerebellar interneurons may perform this role ^72^. Moreover, recent observations suggest that the cerebellar-driven thalamus enables context-dependent responses in the cortex for movement initiation ^51,56^ and cognitive tasks ^73^. Indeed, our work suggests that fast context-switching is easier to incorporate in the relatively simple, divergent and rapidly learnable feedforward architecture of the cerebellum compared to the highly intricate cortical RNNs with weak plasticity.

There are a number of other well described cerebellar properties that would be of interest to study in the context of our framework. For example, incorporating the pontine, cerebellar and thalamic nuclei as intermediate filters ^74,75^, enforcing sparse mossy fibre connectivity ^34,35,76^, and considering synaptic plasticity driven primarily by long-term depression ^32^, are all likely to offer important biological and computational insights.

A unifying model of the cortico-cerebellar loop, and indeed the cerebellum itself, must extend to non-motor tasks. Recent task-based fMRI studies have revealed functional diversity of the cerebellar cortex across a range of cognitive functions ^23^. Our model inherently implies a high degree of heterogeneity – it suggests that different modules would be required to drive different parts of the cortex that in turn underlie different cognitive functions. In this study we modeled recent behavioural and optogenetic experimental observations ^25,26^ which directly implicate the cerebellum in supporting cortical dynamics during evidence accumulation and delayed association tasks (Figs. 5 and 6). In particular, our results show that cortico-cerebellar interactions are enough to learn tasks with highly sparse teaching signals (i.e. only at the end of the task). Furthermore, the model predicts that the cerebellar influence becomes most pronounced during longer task durations (typically in the order of seconds). This phenomenon is attributed to both the preservation of task-specific dynamics through the cortico-cerebellar loop and the cerebellum’s intrinsic capacity, which is enhanced by its extensive hidden granular layer, to disentangle task-specific information from overlapping cortical dynamics. Significantly, we can best capture experimental observations in conditions in which RNN plasticity is limited, making the prediction that the cerebellum is the primary site of learning for these tasks. This provides an alternative to the commonly assumed view that cortical areas are optimised for specific tasks ^6–8^.

In our model the cerebellum drives cortical dynamics based on prediction error signals that depend on the desired task outcome. In the case of the working memory tasks and in line with the experimental task setup, the desired outcome can be interpreted as a reward signal. Therefore, from this perspective, the cerebellum learns to predict future rewarding events. This is consistent with the growing literature showing that the cerebellum encodes reward-related signals ^61,77,78^ and receives projections from the reward system ^79^. However, it remains to be tested exactly how the reward-predictive representations developed by our model compare to those found experimentally.

Here we have also introduced a theory of cerebello-cortical task consolidation. Our theory suggests that cerebellar and cortical learning may operate at different timescales: after an initial fast stage of learning driven by the cerebellum, a period of consolidation ensue in which the cortex gradually acquires task-specific knowledge encoded in the cerebellum (Fig. 7). This view of systems task consolidation is in line with growing experimental evidence suggesting an important role of cerebellar-to-cortical task consolidation ^69,80,81^. For example, Xu et al. ^69^ have observed similar replay-like cerebellar-to-cortical task-specific neuronal dynamics in awake and sleep. Such combination of fast and gradual learning is reminiscent of recent experimental results which suggest significantly faster timescales of plasticity in the hippocampus compared to the prefrontal cortex during a cognitive task ^82^. Moreover, the consolidation period can be related to the idea that a task-optimised cerebellum can be utilised as a cortical teacher ^30,31^. It is in principle possible for cerebellar-thalamo-cortical projections to support this dual role of the cerebellum as both a driver and teacher of cortical states. Indeed, anatomical evidence suggests that this could occur by providing “driving” and “teaching” input to basal and apical dendrites of cortical pyramidal cells, respectively ^83^.

Our work highlights commonalities of cortico-cerebellar interactions in motor and cognitive tasks alike. However, it also suggests interesting differences. The first marked distinction relates to the increased significance of cerebellar-to-cortical (input) plasticity during pure working memory (Fig. 6). This is in line with recent experimental evidence showing stronger plasticity at higher-order thalamo-cortical pathways ^40^. Indeed, because of the need to sustain information during the delay period without sensory or teaching input, it is advantageous for the network to encode a point attractor-like state (see Supplementary Fig. S17, left). Cerebello-cortical plasticity ^39,40^ may thus enable greater *controllability* of cerebellar feedback to push the network to these states during working memory tasks, but less so in motor-based tasks ^62^ (Supplementary Fig. S15).

Related to the point above, the second difference we highlight is about cerebello-cortical consolidation being more readily achieved when in the presence of networks with stable dynamics (cf. Fig. 7 and Supplementary Fig. S18). We speculate that unstable network dynamics make cerebellar-to-cortical consolidation less reliable. Therefore, we predict that while cerebellar-to-cortical systems consolidation might be possible for near perfected tasks which involve discrete stable representations (e.g. working memory tasks), for tasks which are not yet fully learned, or which require faster, more dynamic responses (as often required in the motor domain), cerebellar control is likely to be required throughout life.

To conclude, our work suggests that while the cortex encodes a stable model of the world, it is the cerebellum that allows for quick and flexible adaptation to new environmental conditions. This new cerebellar-guided knowledge can then be gradually consolidated in the cortex.

## Acknowledgements

We would like to thank the Neural & Machine Learning group, Ellen Boven, Aparna Suvrathan, James M Shine, Paul Dodson, Everton Agnes, Laureline Logiaco, Jake Stroud, James Bennett and Heike Stein for useful feedback. J.P. was funded by a EPSRC Doctoral Training Partnership award (EP/R513179/1), P.C. by the Wellcome Trust (209453/Z/17/Z) and R.P.C. by the Medical Research Council (MR/X006107/1), BBSRC (BB/X013340/1) and a ERC-UKRI Frontier Research Guarantee Grant (EP/Y027841/1). This work made use of the HPC system Blue Pebble at the University of Bristol, UK. We would like to thank Dr Stewart for a donation that supported the purchase of GPU nodes embedded in the Blue Pebble HPC system.

## Author contributions

J.P. developed computational framework with guidance from R.P.C. J.P. performed all numerical and analytical work. J.P. and R.P.C. wrote the manuscript, with contributions from P.C. R.P.C supervised the project.

## Methods

### Model architecture and training

**Table S1.** Dynamics of the different model variants, where **h**_*t*_ is the cortical RNN state, **z**_*t*_ the readout and **c**_*t*_ cerebellar feedback. For the experiments presented here we set *f* = tanh and *𝒞* is the cerebellar feedforward network with one hidden layer, *𝒞* (*f* (**h**)) = *W*_PF_*f*^*𝒞*^ (*W*_MF_*f* (**h**)). *W*_*hh*_, RNN recurrent weights; *W*_*ih*_, stimulus-to-RNN weights; *W*_rdt_, (cortical) readout weights; *W*_*𝒞h*_, cerebellar-to-RNN weights; *W*_MF_, cerebellar mossy fibre weights; *W*_PF_, cerebellar parallel fibre weights; *f* ^*𝒞*^ set as ReLU.

|  | No feedback | Readout feedback | Cerebellar feedback | No feedback<br>(cerebellar readout) |
| --- | --- | --- | --- | --- |
| $\mathbf{h}_t$ | $\alpha \mathbf{h}_{t-1} + W_{hh} f(\mathbf{h}_{t-1}) + W_{ih} \mathbf{x}_t$ | $\alpha \mathbf{h}_{t-1} + W_{hh} f(\mathbf{h}_{t-1}) + W_{ih} \mathbf{x}_t + W_{zh} \mathbf{z}_t$ | $\alpha \mathbf{h}_{t-1} + W_{hh} f(\mathbf{h}_{t-1}) + W_{ih} \mathbf{x}_t + W_{ch} \mathbf{c}_t$ | $\alpha \mathbf{h}_{t-1} + W_{hh} f(\mathbf{h}_{t-1}) + W_{ih} \mathbf{x}_t$ |
| $\mathbf{z}_t$ | $W_{rdt} f(\mathbf{h}_t)$ | $W_{rdt} f(\mathbf{h}_t)$ | $W_{rdt} f(\mathbf{h}_t)$ | $\mathcal{C}(f(\mathbf{h}_t))$ |
| $\mathbf{c}_t$ | NA | NA | $\mathcal{C}(f(\mathbf{h}_{t-1}))$ | NA |

The complete dynamics of each model architecture that we consider (Supplementary Fig. S1; no feedback, readout feedback, cerebellar feedback, no feedback with cerebellar readout) are given in Table S1. In all of our simulations we use a recurrent neural network (RNN) with 50 time-discrete units (see section below).

Unless otherwise stated, the feedforward cerebellar network contains a single hidden layer with 1000 units (granule cells), but other hidden layer sizes are also considered (Figs. 2D and 4E). This yields a divergence from the cortical RNN to the cerebellar granular layer of 50:1000 = 1:20. The cerebellar output layer, which we interpret as Purkinje cells, on the other hand, mirrors the desired task outcome and is therefore of significantly lower dimensionality (3 in evidence accumulation task and 2 in all other tasks).

For each task simulation, network parameters are initialised as follows. The RNN input, recurrent and cerebellar feedback weights *W*_*ih*_, *W*_*hh*_, *W*_*𝒞h*_ are drawn from a uniform distribution *W* ^init^ ∼ *𝒰* (*−a, a*) where 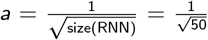. The readout weights *W*_rdt_ and cerebellar weights, *W*_MF_, *W*_PF_, are initialised according to *𝒰*(*−b*_*k*_, *b*_*k*_) where *b*_*k*_ denotes the “kaiming bound” He et al. ^84^ (slope 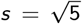). The biases of the cortical readout are drawn from 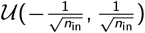, where *n*_in_ denotes the input size of the layer. In line with existing models of cortical networks ^16^, in our model we do not obey Dale’s law and use a tanh activation function. In future work it would be of interest to test a variant of our model with explicit excitatory and inhibitory cortical populations. We conducted each task simulation with 5 random seeds for initialisation.

During the learning of a task model parameters are updated using gradient descent from the task error signal *E* = ∑_*t*_ *E*_*t*_ with respect to to the model parameters (see section below). For each dataset each training session covers 1000 random examples, presented to the model in batch sizes of 10 which we call a “trial”. The test set (used after training) also covers 1000 randomly generated examples. When analysing the learned network dynamics (e.g. model output with and without cerebellar ablation) the model with the best validation error during training was selected. An ADAM optimiser ^85^ was used with initial learning rate *η* = 0.001 for the RNN (when plastic), readout and cerebellar network, except for the delayed association task for which we found an RNN learning rate of *η* = 0.0025 to provide more stable learning. The different plasticity constraints of the entire model - termed “fixed RNN”, “input plastic”, and “fully plastic” - are defined with respect to the cortical parameters of Eq. 1 as follows. For the fixed RNN case, only the cortical readout weights *W*_rdt_ are learned. For the input plastic case, RNN input weights and *W*_*ih*_ and *W*_*𝒞h*_ are also learned. Finally, for the fully plastic case, the recurrent weight *W*_*hh*_ is also learned. In all of these cases the cerebellar “parallel fibres” *W*_PF_ are learned, whilst the “mossy fibres” *W*_MF_ remain constant, in line with mossy fibres synapses being (relatively) stable ^33,86^.

In each of the considered tasks we report the change in error during and after training as a result of cerebellar feedback (Figs. 2L,4I,5H,6I,). The change in error during training is computed as the average difference in training error between the cerebellar feedback and no feedback models. The change in error after training is computed as the average difference in test error between a trained cerebellar feedback model, and a trained cerebellar feedback model subject to cerebellar ablation. As in the main results this cerebellar ablation after training may be transient. In particular, for the line drawing and digit drawing tasks we consider transient ablation during the middle period of the task, for the delayed association task we consider transient ablation as Fig. 6D-G, and for the evidence accumulation task we consider full cerebellar ablation.

#### Continuous dynamics of RNN model

A continuous version of our RNN can be expressed as

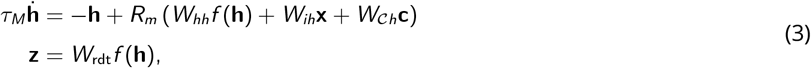

where *τ*_*M*_ is the membrane time constant (not to be confused with the cerebellar time window *τ*), *R*_*m*_ is the membrane resistance, and *f* is the rate-based non-linearity which we set as *f* = tanh. Discretising Eqs. 3 with timesteps of Δ*t* yields equations in Table S1, where 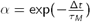. Note that as in ^38^ we ignore the (1 *− α*)*R*_*m*_. This simplifies notation and has no effect on dynamics if model weights are scaled accordingly. In general we use *τ*_*M*_ *≈* 20ms and Δ*t* = 50ms for the drawing tasks (Figs. 2,3 and 4) and a higher *τ*_*M*_ *≈* 90ms with Δ*t* = 200ms for the cognitive tasks (Figs. 5,6 and 7) in line with ^6^). In both cases this gives us a cortical internal memory *α* = 0.1.

### Cortical and cerebellar learning rules

When the desired task outcome **y**_*t*_ is provided the associated error is computed as *E*_*t*_ = *ℰ*(**z**_*t*_, **y**_*t*_) for the cortical network and 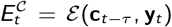 for the cerebellar network, where *E* denotes the task error function (mean squared error and cross-entropy loss for regression and classification tasks respectively) and *τ* is the cerebellar time window. The error gradients for the readout and cerebellar weights *W*_rdt_, *W*_PF_ can then be obtained locally with a simple delta-rule on the gradient of the error signal. That is,

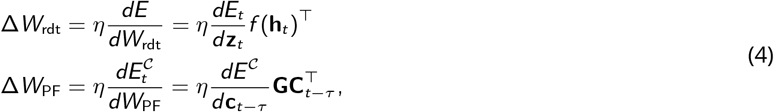

where *η* denotes the learning rate of the cortico-cerebellar network and **GC** denotes the hidden granule cell activity of the cerebellar network which is computed as **GC**_*t*_ = *f* ^*𝒞*^ (*W*_MF_*f* (**h**_*t−*1_)) (cf. Eq. 2).

For the input/recurrent weights *W*_*ih*_, *W*_*𝒞h*_, *W*_*hh*_ - when plastic - obtaining error gradients is more difficult as temporal dependencies need to be considered. To improve biological feasibility in this work we avoid backpropagation through time (BPTT) and instead use the eprop algorithm ^38^. Details can be found in ^38^, but the main idea is that BPTT can be approximated with a mixture of locally computed synaptic eligibility traces and current learning signal. Specifically, the error gradient for a given synapse *w*_*ji*_ from neuron *i* to *j* is computed as

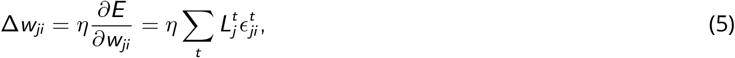

where for ease of notation we now use the superscript to denote timestep *t* and 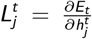 is the neuron *j* learning signal (obtained by one-step backpropagation through space except for the cerebellar readout architecture in Supplementary Fig. S1D). 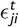 is the synaptic eligibility trace of *w*_*ji*_ which is computed as defined recursively by

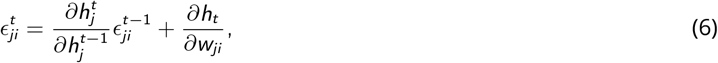

where 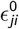 is initialised as zero. Note that the terms in Eq. 6 are locally available to the synapse. In the case of our network dynamics (Eq. 1), the eligibility trace is simply defined by 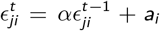, where *a*_*i*_ is the activation of the presynaptic neuron *i* (e.g. tanh(*h*_*i*_) or *c*_*i*_).

For all weights, the error gradients are accumulated across multiple examples (i.e. batch update) and timesteps before the weights themselves are updated.

#### Learning rules for cerebellar-to-cortical consolidation

A period of “consolidation” is considered for the trained models of the delayed association task (Fig. 7 and Supplementary Fig. S18). During this period the model is presented with further trials (batch size 10) of training data but without their associated targets. The forward dynamics of the model then run as normal (Eq. 1) but now we use a consolidation learning rule for the RNN weights. We consider both an optimal learning rule which uses the least-squares algorithm and also a simple biological learning rule.

We first present the optimal consolidation learning rule, since this motivates the biological rule. We want to change the recurrent (cortico-cortical) input to match the cerebellar-cortico input over the task. To this end we concatenate the time-dependent RNN activities 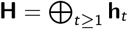 and cerebellar output activities 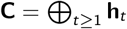, where ⊕ denotes vector concatenation. We then set the change in recurrent weight Δ^cons^*W*_*hh*_ with 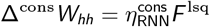 where 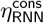 consolidation learning rate and *F* ^lsq^ is the least-squares solution

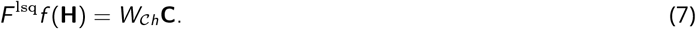

At the same time the cerebellar-cortical weights *W*_*𝒞h*_ decay according to

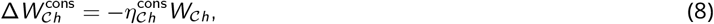

where 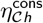 is the rate of cerebellar-cortical decay. In the experiments shown we select 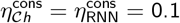.

For the biological learning rule, the cerebellar-cortical weight decays as in Eq. 8 but now the RNN weights are updated according to the ratio of cerebellar feedback against the whole population activity. That is, for the recurrent weight from neuron *i* to neuron *j* we have

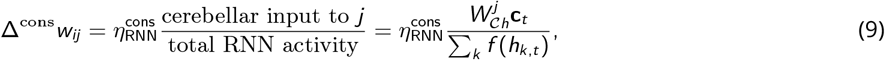

for arbitrary timestep *t* and where 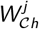 denotes the *j*th row of the cerebellar-cortical weight *W*_*𝒞h*_.

To demonstrate that Eq. 9 leads to changes in cortico-cortico input which are proportional to the cerebellar-cortical input, we see that the change in recurrent input to a given RNN neuron *j* at time *t* becomes

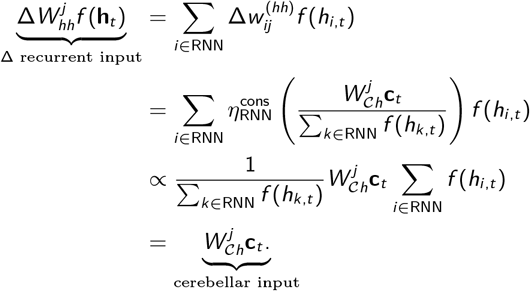

That is, we recover a solution (up to proportionality) to Eq. 7. For this biological learning rule, to improve network stability, we found it beneficial to increase the RNN consolidation learning rate such that 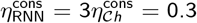 (where Δ^cons^*w*_*ij*_ is accumulated over the whole sequence). This explains the initially faster learning (over the first few trials) for the biological learning rule (Fig. 7F).

For this consolidation learning period a learning optimiser is not used (i.e. ADAM is not used). Note that these consolidation learning rules do not require information about the desired task outcome (i.e. target) and are in that sense unsupervised.

#### Demixed principal component analysis

To study the response dynamics specific to task variables in the delayed association task (Fig. 6) we perform demixed principal component analysis (dPCA) ^63^. dPCA extracts low-dimensional components that explain maximum population variance constrained by task-specific variables. As a result we obtain principal components that are specific to task variables; in this case the task variable of interest is animal/model choice. The neural data we provide as input to dPCA is a three-dimensional array (*n, s, t*) with each dimension representing average neuronal activity (concatenated across animals/seeds), choice identity and time, respectively. dPCA is applied to the model representations (after learning) and neural data acquired in ^25^.

### Task details

#### 1. Line drawing task

For the line drawing task, the model has to transform one of six possible 10-dimensional binary inputs **x** ∈ [0, 1]^10^ at timestep 1 into an associative “go” 2-dimensional line **y**^line^ (for five of the inputs) or a “no-go” stay at the origin (for one of the inputs). The starting point for each line is the origin, and the endpoints of each line are evenly spaced on the edge of the unit circle (see Fig. 2A, black dashed line). The model learns to draw the line over 20 discrete timesteps, with the intermediate target points spaced evenly, i.e. for a line with endpoint *y*_end_ we have 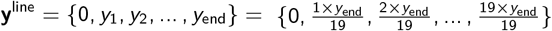.

For the stimulus timestep (timestep 1) as well as the remaining 19 timesteps, the model receives (through its *W*_*ih*_ connection) zero-mean Gaussian noise *ξ* ∼ *N* (0; 0.1^2^). Model errors are computed as the mean-squared error to the target response. Unless otherwise stated a cerebellar time window *τ* = 3 timesteps (*≈* 150ms when *α* = 0.1) is used. The prediction error across time delay *t*_0_ between cortical output and cerebellar (or cortical) output (Fig. 2E) is computed as the cue/time average 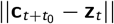, where ||.|| is the Euclidean norm.

To analyse the effects of cerebellar ablation we consider partial cerebellar ablation at the start, middle, and end of the sequence (Fig. 2h-k and Supplementary Fig. S5). The specific time windows of these ablation periods are timesteps [1-6, 8-13, 15-20] (inclusive), respectively.

##### Curl-ield variant

Once the models of the line drawing task are trained, we tested whether they could re-translate the same external inputs to a curl-field variant of the task (see e.g. ^54^). For this we selected models with cortical internal memory *α* = 0.5, since we found this resulted in faster learning which was comparable to the presented experimental data ^54^, but we find *α* = 0.1 (as presented in Fig. 2) also learns but more slowly. Switching and learning this curl-field new task “context” involved retraining the models to new desired outcomes (central grey curves in Fig. 3C).

Specifically, the curl-field target responses have the same end-point for each line (or same “no-go” zero cue), but intermediate target points now form a semi-ellipse between the origin and the respective end-point. Given the desired endpoint 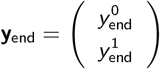, this can be parameterised by

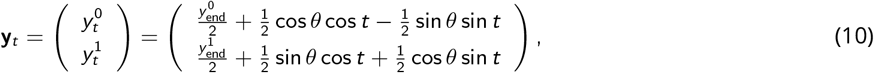

where 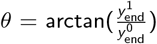 is the angle to the end point and *t* runs uniformly between 0 and *π* (or, for direction towards (*x*_end_, *y*_end_) as in our experiments, from *π* to 2*π*).

To test how context-dependent cerebellar processing could enable rapid task switching, we considered the extent to which parallel fibre (PF) weights are shared across task contexts. In particular, we label the percentage of PFs used for each context as the PF task overlap. For example, if the PF task overlap is 25%, then 25% of the PFs used for cerebellar processing apply to both task contexts, whilst 75% specifically apply (and are trained) to the current context. Before learning, the PFs which are not shared (i.e. only apply to the curl-field context) are initialised randomly as in the original line-drawing task.

##### Neuronal activity and covariance during task switching

The change in activities and change in covariances (Fig. 3D-F and Supplementary Fig. S7) are computed as in ^58^. We record the RNN time-dependent activities (post non-linearity) given 1000 input examples in multiple periods: task 1 baseline, task 2 and task 1 switching (Fig. 3A). For the latter two periods these are recorded at their respective end, whilst we take two samples of the baseline period at its start and end. The change in activity between any two periods *P*1 and *P*2 is the average change in activity for a given neuron *i*, which is given by

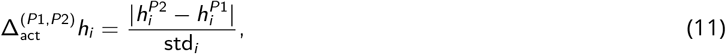

where 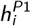, 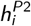 are the time-varying input-dependent activities of neuron *i* for periods *P*1, *P*2 respectively, and std_*i*_ is the standard deviation of that neuron in the start of the task 1 baseline period. Here |.| denotes the average (absolute) difference in activity across timesteps and input examples.

For each period, we also compute the covariance matrix of the RNN population. The change in covariance between two sessions 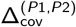 is then computed as 1 minus the Pearson correlation between their respective covariance matrices ^58^.

For the task 2 and task 1 switching periods we report changes with respect to the start of the task 1 baseline period. To account for natural variability in the network and better compare to the neural data in ^54^, we normalise the changes by taking away the changes observed within the baseline period itself. For example, the change in covariance in the task 2 period is 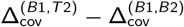, where *B*1, *B*2, *T* 2 are the start of the task 1 baseline, end of task 1 baseline, and (end of) task 2 respectively. We apply the same normalisation to the reported experimental changes in the monkey M1 and PmD ^54^; this normalisation leads to (average) near-zero change for the M1 activity and PmD (Fig. 3F).

The number of training trials for training in task 2 shown in Fig. 3A (500 trials) leads to good, but not perfect, performance. To demonstrate that the models can eventually perform task 2 to a close to perfect standard, the model outputs presented in Fig. 3C underwent 1000 trials of training.

#### 2. Digit drawing task

For the digit drawing task the inputs are the same as the 10-dimensional binary vectors used in the line drawing task, except now the model must draw an associative digit over 20 timesteps instead of line (Fig. 4A). The targets **y**^digit^ are constructed manually within the space [0, 1]^2^ and resemble the digits from 0 to 5 (inclusive). For exact implementation refer to the provided code (see below).

For the standard model with cerebellar feedback a cerebellar time window *τ* = 3 timesteps (*≈* 150ms when *α* = 0.1) is generally used. For the model using cerebellar feedback with a temporal basis, we model the cerebellum with a range of time windows, i.e. *τ* = {*τ*_*i*_}_*i*_ for some distinct *τ*_*i*_ ≥ 0ms. In this task we consider 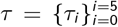 with *τ*_*i*_ = *i* timesteps (i.e. 0-250ms), so that the final cerebellar output is a concatenation of task predictions which span over the proceeding 250ms period. Explicitly, after training we have cerebellar feedback, 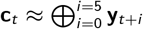, where ⊕ denotes vector concatenation.

Zero-mean Gaussian noise *ξ* ∼ *𝒩* (0; 0.1^2^) is added to the input at each timestep. Model errors are computed as the mean-squared error to the target response.

To analyse the effects of cerebellar ablation we consider the same partial cerebellar ablation periods as in the line-drawing task. That is, we consider cerebellar ablation at the start, middle, and end of the sequence (Fig. 4 and Supplementary Fig. S9), which correspond to timesteps [1-6, 8-13, 15-20] (inclusive), respectively.

#### 3. Evidence accumulation task

In the evidence accumulation task the model receives 2-dimensional binary inputs (i.e. **x** ∈ [0, 1]^2^) over a presentation period of *T* ^pres^ = 45 timesteps. A non-zero input can occur for at most one of the two dimensions; that is, *x*_*t*_ ∈ {(1 0)^⊤^, (0 1)^⊤^, (0 0)^⊤^}, where the rate of zero inputs *x*_*t*_ = (0 0)^⊤^ defines the sparsity of input *ρ* (*ρ* = 0.7 in our simulations). After this presentation of input there is then a delay period of *T* ^del^ = 5 timesteps after which the model must classify at which dimension more non-zero input was received (or whether the number at each dimension was the same). That is, the desired outcome *y* takes one of three values which respectively correspond to more input in the first dimension, more input in the second dimension, or the same. This task resembles the experimental structure of ^26^, in which mice were trained to select the side of their whiskers which received more air puffs.

Zero-mean Gaussian noise *ξ* ∼ *N* (0; 0.1^2^) is added to the input at each timestep. Model errors are defined by the cross-entropy loss to the target response.. Model “belief” (Figs 5D and S11) is defined as the model probability (obtained by applying a softmax on the readout) of the correct classification. Unless otherwise stated a cerebellar time window *τ* = 3 timesteps (*≈* 600ms when *α* = 0.1) is used. For both readout and cerebellar feedback models, we apply a softmax operation to the feedback returned to the RNN so as to bound its values between 0 and 1.

To analyse the effects of cerebellar ablation we consider full cerebellar ablation (for the entire sequence 1-50; see Fig. 5D and Supplementary Fig. S11A-C, left) and also partial periods of ablation: at the start, middle, and end of the sequence (Fig. 5E,F and Supplementary Fig. S11A-C, right). The specific time windows of these partial ablation periods are timesteps [1-15, 15-30, 30-45] (inclusive), respectively. To improve readability of our results, the mean error presented in the training curves for this task is smoothed using a Savitzky-Golay filter with window length 25 and polynomial order 3.

To compute the dependence of model choice on inputs over different temporal bins (Fig. 5F), we follow the method in ^26^. In particular, we divide the presentation period evenly into 3 time windows - [1-15, 16-30, 31-45] - and fit the model choice according to a logistic regression model

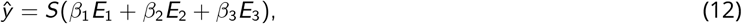

where *ŷ* denotes the predicted model choice probability, *S* is the sigmoid logistic function, *E*_*i*_ = #*R*_*i*_ *−* #*L*_*i*_ is the different in the total number of ‘right’ and ‘left’ inputs in window *i*, and *β*_*i*_ is the respective weight on that window. *ŷ* is fitted to minimise the negative log likelihood of the observed model decisions. We present the normalised weights of each window 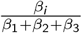.

##### History-centric cases

In line with ^26^, we observe cerebellar ablation to be particularly detrimental to input examples for which correct classification would depend on adequately maintaining past inputs (Fig. 5E,F and Supplementary Fig. S11), which we refer to as “history-centric” examples. We define an input example as being history-centric if exposure only to the final third of the input sequence would lead strictly to the wrong answer. That is, examples (**x**, *y*) such that the “final-third target” 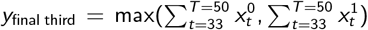 is not equal to the desired outcome 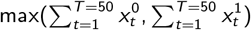.

##### Sub-second task lengths

To identify whether dependency on cerebellar feedback holds for shorter timescales, we consider cue presentation periods from 0.1 *−* 1s (Fig. 5G). For these simulations there is no delay period and the sparsity of input is *ρ* = 0.5. We apply a finer time discretisation so that Δ*t* = 10ms; we redefine the cortical internal memory *α* and rescale the network parameters accordingly. The cerebellar network is trained with a time window *τ* = 3 timesteps in each case.

#### 4. Delayed association task

In the delayed association task the model must associate one of two 10-dimensional binary inputs at timestep 1 to a desired binary response *y* at timestep *T*, where *T* is the sequence length or “delay” period ^25^. We select *T* = 15 timesteps but also consider other lengths (Fig. 6J). The task error (as presented in the main text) is defined at the end of the sequence. For stability, we train the network output 5 timesteps from the end of the sequence (timestep 10 onwards when *T* = 15).

Zero-mean Gaussian noise *ξ* ∼ *𝒩* (0; 0.1^2^) is added to the input at each timestep. Model errors are defined by the cross-entropy loss to the target response. Model “selectivity” is defined as the model output (readout) at the dimension of the correct classification (prior to the softmax operation). Unless otherwise stated a cerebellar time window *τ* = 3 timesteps (*≈* 600ms when *α* = 0.1) is used. For both readout and cerebellar feedback models, we apply a softmax operation to the feedback returned to the RNN so as to bound its values between 0 and 1.

To analyse the effects of cerebellar ablation we consider cerebellar ablation within a particular time window between timesteps 8-12 (inclusive) which approximately mirrors the timings in ^25^ (Fig. 6D,E and Supplementary Fig. S17) and also partial ablation periods during the start, middle, and end of the sequence (Fig. 6F). The specific time windows of these partial ablation periods are timesteps [1-5, 6-10, 11-15], respectively. To improve readability of our results, the mean error presented in the training curves for this task is smoothed using a Savitzky-Golay filter with window length 25 and polynomial order 3.

For this task we consider how the model evolves during a consolidation period (Fig. 7). At the end of each consolidation trial we observe the model error (Fig. 7B), activity (Fig. 7E) and recurrent input (Fig. 7F) over a test set of 1000 randomly generated examples. The activity here is the concatenation of activity in the cortical RNN and the hidden layer of the cerebellar network (over all examples and timesteps). We compute the cosine similarity between these activities and the initial activities prior to consolidation; for comparison we also the cosine similarity been the initial activities and a shuffled version of the initial activities (averaged over 100 samples). To analyse how the recurrent input changes we proceed as follows. At each timestep we consider the cortical RNN state **h** and cerebellar feedback **c**. We then compute the cosine similarity between *W*_*hh*_**h** and 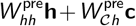, where 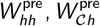 are the pre-consolidation RNN weights and cerebellar-cortical weights, respectively.

### Control-theoretic estimation of cerebellar feedback

For the delayed association task we analyse cerebellar-to-cortical input from a control-theoretic point of view. In particular, we quantify the effect of plasticity in the pathway between the cerebellar network and cortical RNN (*W*_*𝒞*_*h*) on cortical activations by estimating the energy cerebellar feedback induces in RNN state space ^62^. This level of energy reflects the potency of feedback onto the RNN: a low energy would reveal a suppressed RNN response, whereas a high energy would reveal an amplified response. We speculated that these two cases would arise from a non-optimised *W*_*𝒞⟨*_ and optimised *W*_*𝒞⟨*_, respectively (Supplementary Fig. S15A).

As per Kao and Hennequin ^62^, we compute the energy of cerebellar feedback through the *controllability Gramian P* associated with RNN dynamics. Informally, *P* describes the “intrinsic manifold” of the RNN and describes the directions in which the RNN is most (or least) likely to visit. Formally, given a direction **v** in state space, the average energy generated along direction *v* is

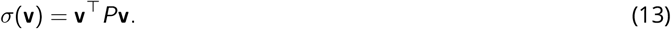

In general, the Gramian matrix *P* is only defined for linear systems. In this work we therefore generalise the notion of controllability for the non-linear RNN dynamics as defined in Eq. 1. Here we use the noise covariance matrix ∑ in its place, which for linear systems is shown to be equivalent to the Gramian, ∑ = *P* ^62^. Explicitly, we compute ∑ as the time-course average covariance of RNN hidden activations *h*_*t*_ under noisy inputs which follow a Wiener process. That is, ∑ = 𝔼_*t*_ [cov(*H*_*t*_)] where 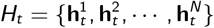 is a set of *N* samples of RNN states which each evolve according to

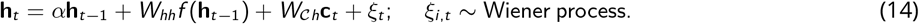

In our experiments we use *N* = 500 samples and simulate Eq. 14. To ignore intrinsic RNN transients that occur at the start of simulation, we discard the RNN states during the first 5 simulation timesteps when computing ∑. The energy generated from cerebellar feedback is then *σ*(**h**^*𝒞*^) = (**h**^*𝒞*^)^⊤^∑**h**^*𝒞*^, where 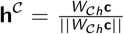 is the normalised direction being driven by the cerebellum in RNN state space. We report the energy generated (during the noise dynamics of Eq. 14) by cerebellar feedback at timestep 10, a time chosen strictly after the initial RNN transient phase (Supplementary Fig. S15B). For comparison we compare this to the energy generated by 100 random sample directions **v** ∼ *𝒩* (0, *I*) where *I* is the identity matrix. To enable greater interpretability we then normalise these energies by its highest possible value max_||**v**||=1_ **v**^⊤^∑**v**; i.e. the input which elicits maximal amplification of RNN dynamics. This value can be computed as **u**^⊤^∑**u** where **u** is the principal eigenvector of ∑.

### Cerebellum decodes low-signal cortical representations

For the delayed association task we discussed the need for a greater number of hidden cerebellar units (granule cells) to achieve good task performance (Fig. 6J). In particular, we find that the number of granule cells (GCs) required is inversely proportional to the *signal-to-noise* (SNR) of the RNN hidden neurons.

To estimate SNR(RNN) in the models for the delayed association task (Fig. 6K, left axis), we suppose that the activity population activity in the RNN can be divided into two components such that *f* (**h**) = *ζ* +*ω*, where *ζ* is a task-dependent component which depends on the current task condition *s* (i.e. left or right stimulus), and *ω* is a task agnostic component which does not depend on *s* (but instead depends on, for example, intrinsic RNN connectivity and noise). The SNR is then defined as the ratio of the variance of these two respective components: 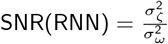

We compute the variance of the task-agnostic component as the (average) variance of the population under the same task stimulus *s*, i.e. 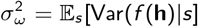. Be equally calculating the total variance 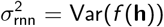, the variance of the task-relevant component is then simply computed as the difference to the total variance, i.e. 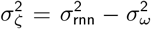. To determine the minimum number of granule cells required to decode the stimulus from the RNN activity (Fig. 6K, right axis), we tested whether the cerebellar network could be trained to successfully discriminate the stimulus after 40 training sessions for varying quantities of granule cells (quantities as described below). The cerebellar network was deemed to successfully decode the stimulus if, for at least 4 of the 5 seeds, the average error during the last 4 training sessions was less than 5%.

## Data and code availability

We used the PyTorch library for all neural network models. The code and respective simulated data used for our experiments is available at https://github.com/neuralml/ccLoops.

## Supplementary Information

**Figure S1.**
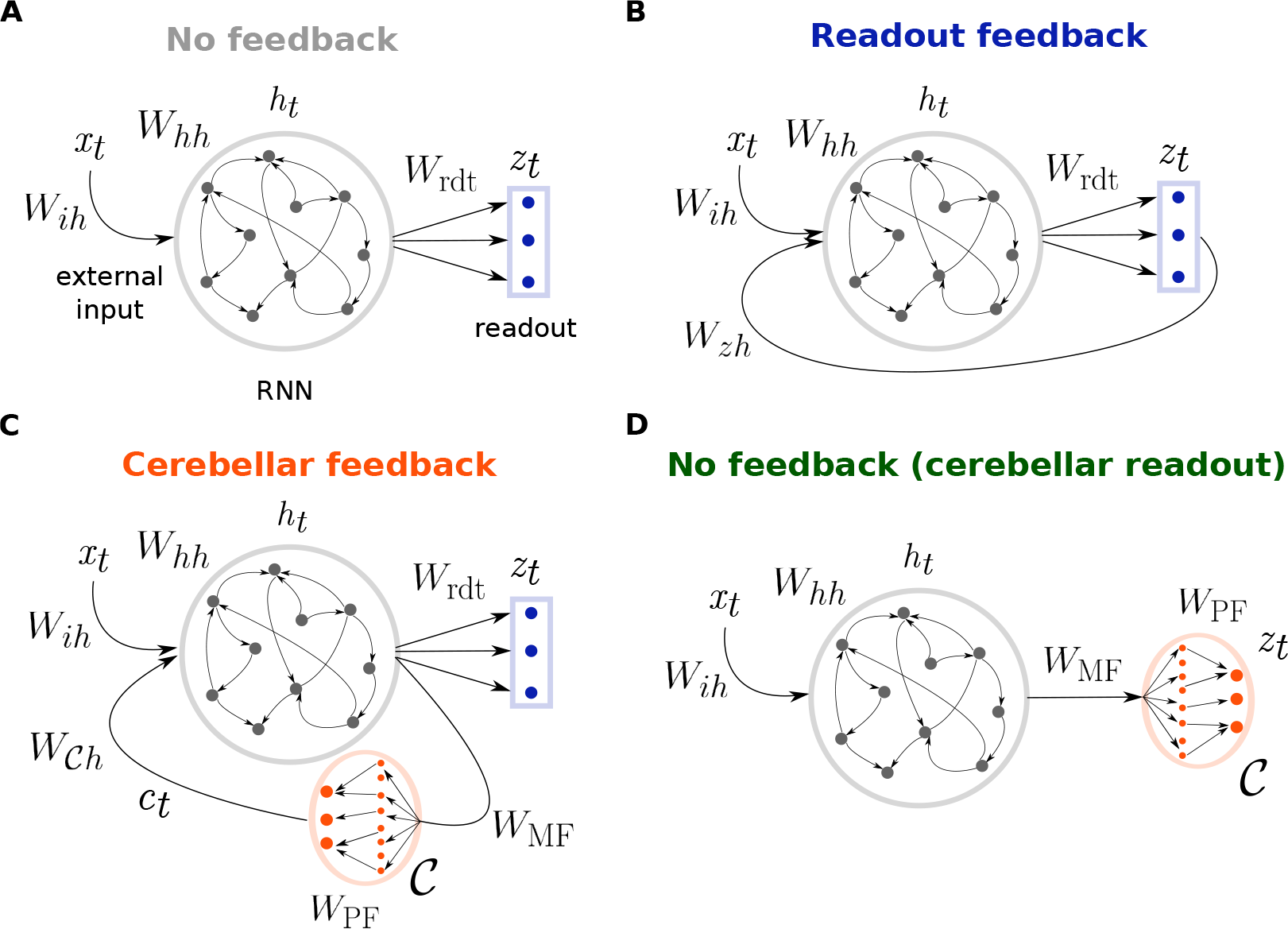
Different model architectures (extension of Fig. 1). (**A**) *No feedback*; temporal input is fed to a cortical RNN (grey) and a linear readout layer (blue) produces the final model output. (**B**) *Readout feedback*; now there is a feedback loop in which the RNN also receives readout predictions as extra input ^14,16^. (**C**) *Cerebellar feedback*; a copy of RNN activity is sent to a distinct but connected cerebellar network *C*, which then returns its predictions back to the RNN as extra input. (**D**) *No feedback with cerebellar readout*; like in C a cerebellar network is attached to the RNN, but now it is used directly as the final readout and there is no “cortico-cerebellar loop”. Model activity and weight vectors are represented with the same notation as Eqs. 1 and 2 (see also Table S1).

**Figure S2.**
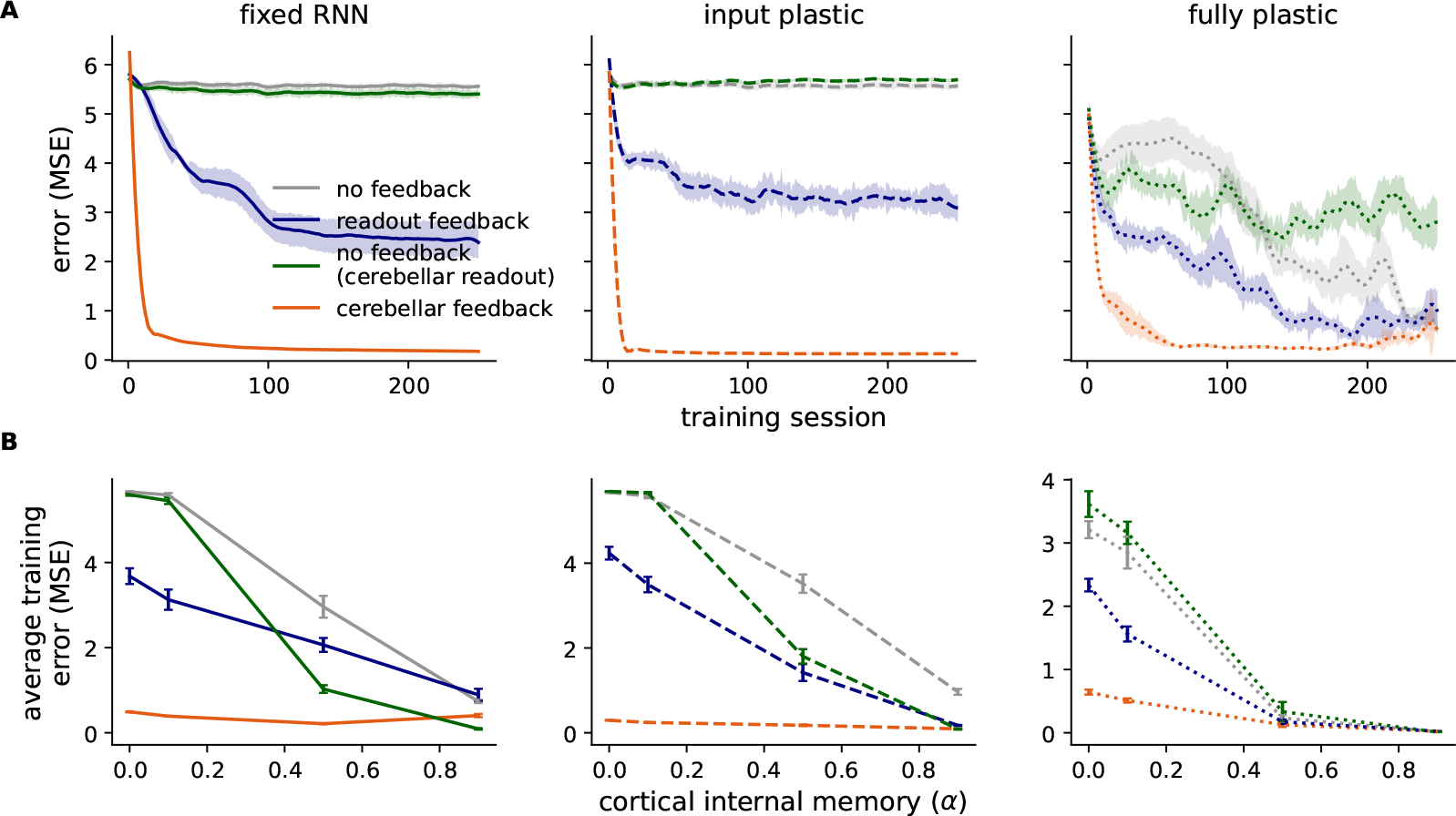
Model learning in the line drawing task. (**A**) Training curves (cortical internal memory *α* = 0.1) for the different models with fixed (left), input plastic (middle) and fully plastic (right) RNN. Green denotes the model where no feedback is applied to the RNN but the readout network (usually linear) now has the same architecture as the cerebellar network. (**B**) Average error over training across different cortical internal memory *α*.

**Figure S3.**
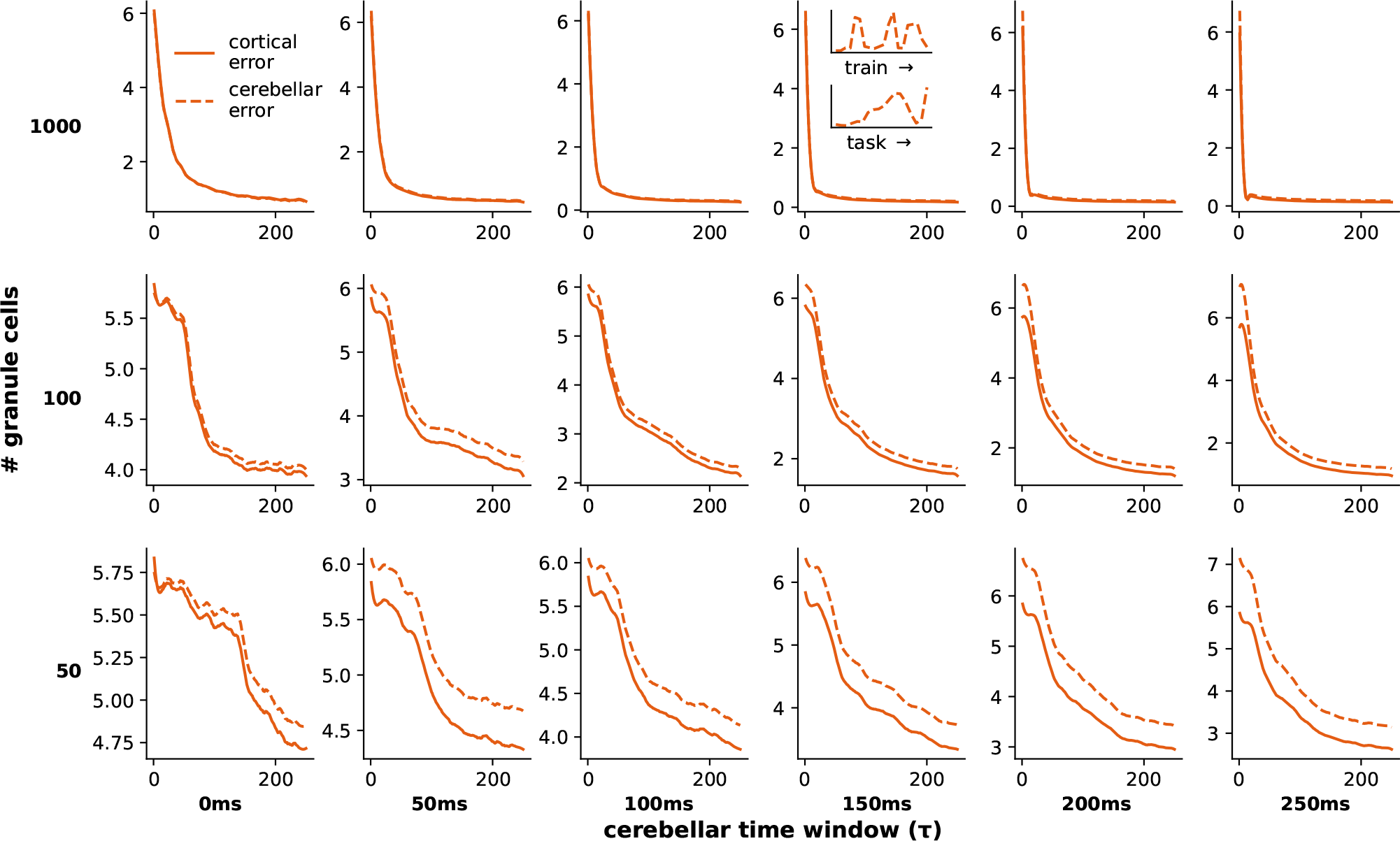
Training curves over different cerebellar parameters. We show the learning curves for the cortical network (solid line) and cerebellar network (dotted line) for the line drawing task. On each miniplot the x-axis represents the training session and y-axis the mean-squared error. The cerebellar error for an example seed is shown in the inset of the model conditions used in the main text (1000 granule cells, time window *τ* = 150*ms*), over different task examples during training (upper) and over time within one task example (lower).

**Figure S4.**
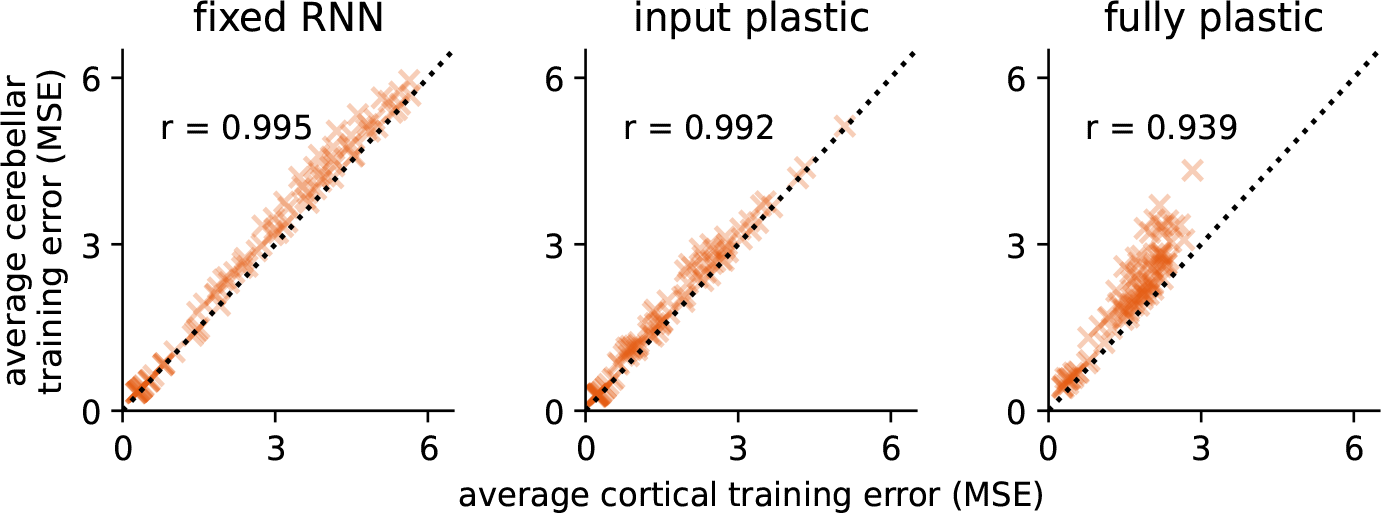
Integrated errors for the cortical network against the cerebellar network for the line drawing task. Each point denotes a specific cerebellar parameter configuration (1 of the 18 in Supplementary Fig. S3) and initialisation seed (1 of 5). *r* denotes the Pearson correlation.

**Figure S5.**
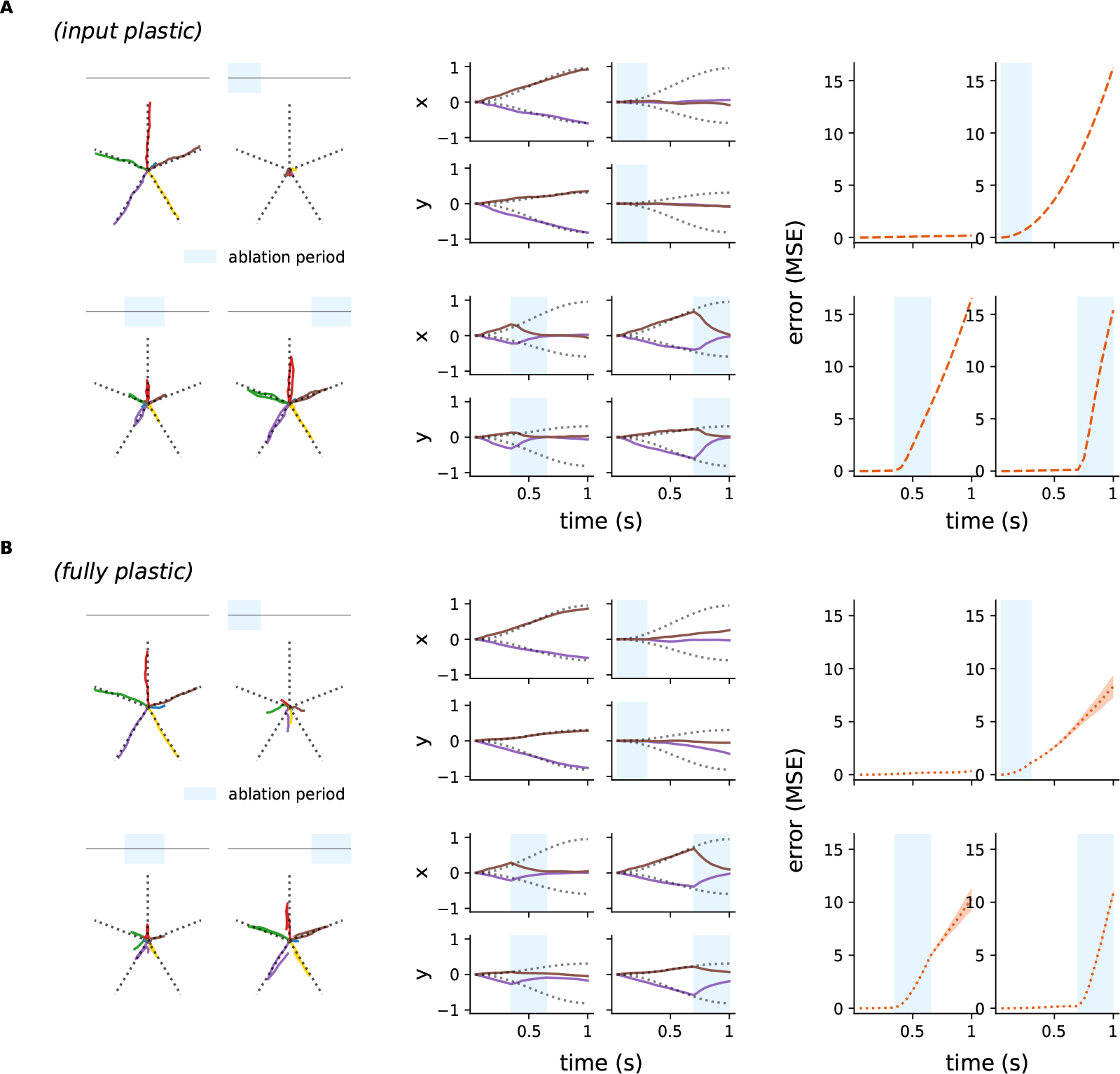
Ablation results for the input and fully plastic RNN. Model output (left, middle) and error (right) for example line drawing input under cerebellar ablation for an (**A**) input plastic and (**B**) fully plastic RNN. For the corresponding fixed RNN case see Fig. 2H-J.

**Figure S6.**
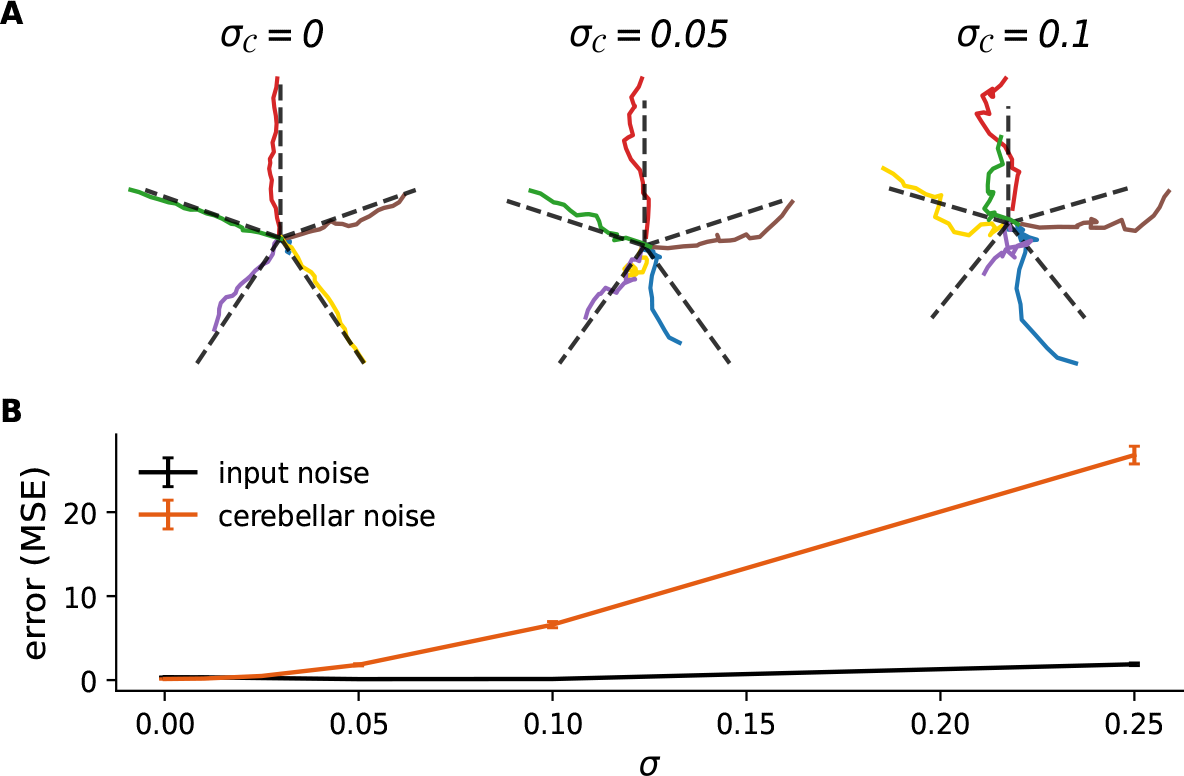
Cerebellar noise induces ataxic-like impairments. (**A**) Output for trained models on the line drawing task under different levels of cerebellar noise 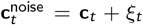 with 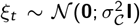; blue output is for “no go” cue (where model is trained to remain at zero). (**B**) Model error under various degrees of input and cerebellar noise.

**Figure S7.**
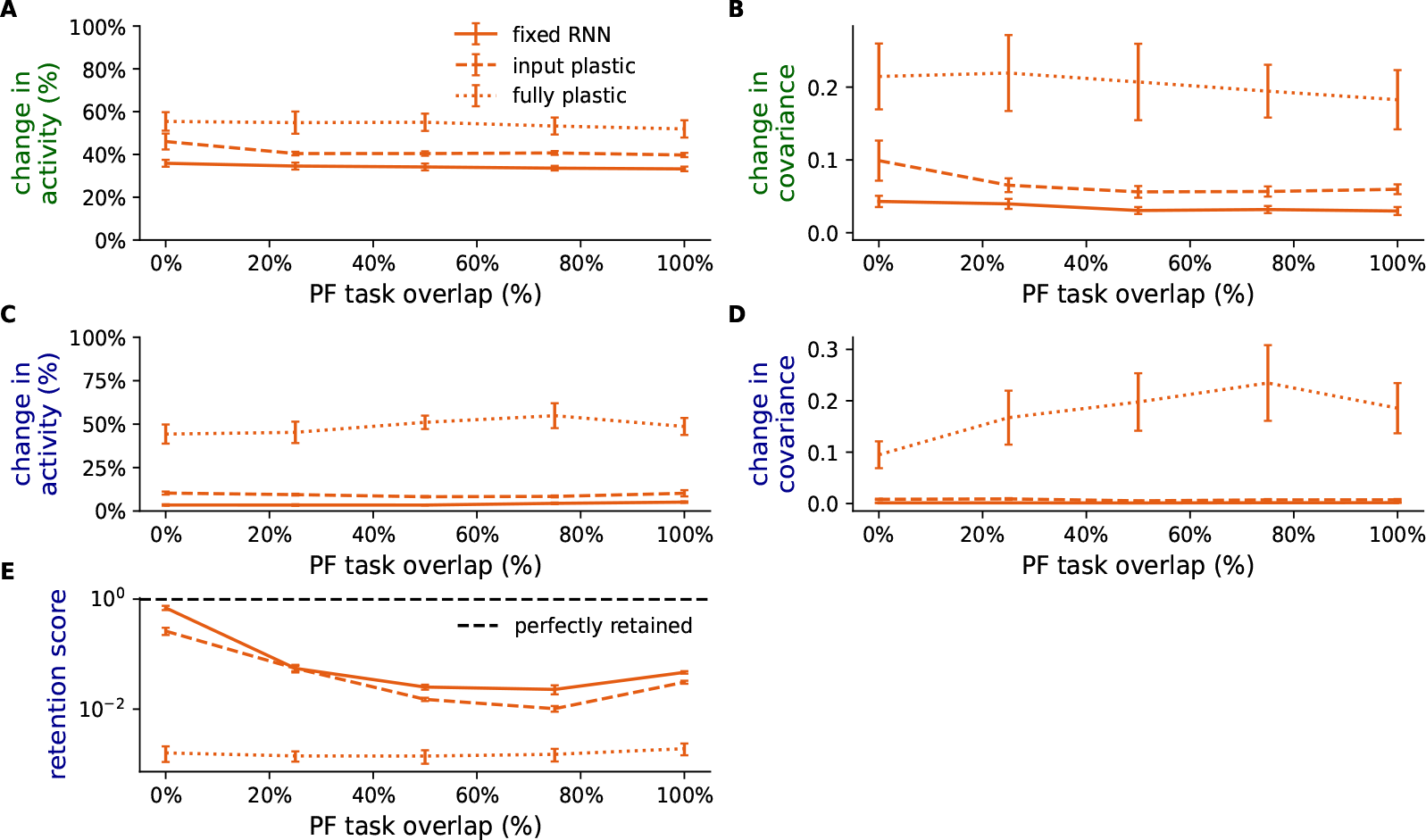
Multi-task learning and switching across different levels of parallel fibre (PF) overlap. (**A**,**B**) Change in (A) activity and (B) covariance in RNN population between the line-drawing task 1 (baseline) and task 2 which is a curl-field variant. (**C, D**) Change in (C) activity and (D) covariance in RNN population between task 1 (baseline) and task 1 (post re-learning) after switching back from task 2. (**E**) Task 1 retention score, which is computed as the error of task 1 during baseline over the error at the first trial after switching back to task 1.

**Figure S8.**
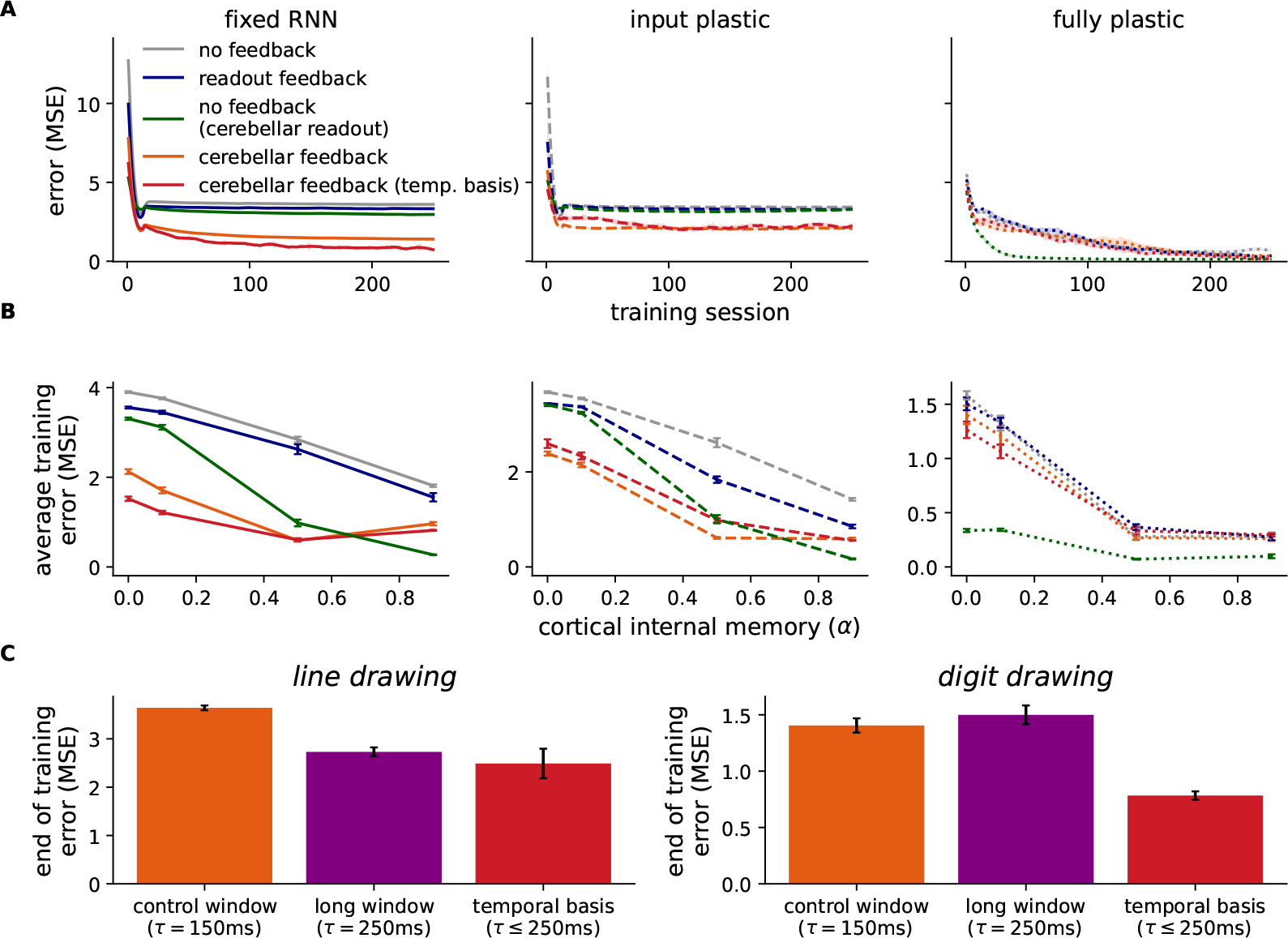
Model learning in the digit drawing task. (**A**) Training curves (cortical internal memory *α* = 0.1) for the different models with a fixed (left), input plastic (middle) and fully plastic (right) RNN plasticity assumptions. Green denotes the model where no feedback is applied to the RNN but the readout network (usually linear) now has the same architecture as the cerebellar network. (**B**) Average error over training across different cortical internal memory *α*. (**C**) Model error (fixed RNN; *α* = 0.1) at the end of training (averaged over last 10 training sessions) for different cerebellar time windows for (left) line drawing task (cf. Fig. 2) and (right) digit drawing task.

**Figure S9.**
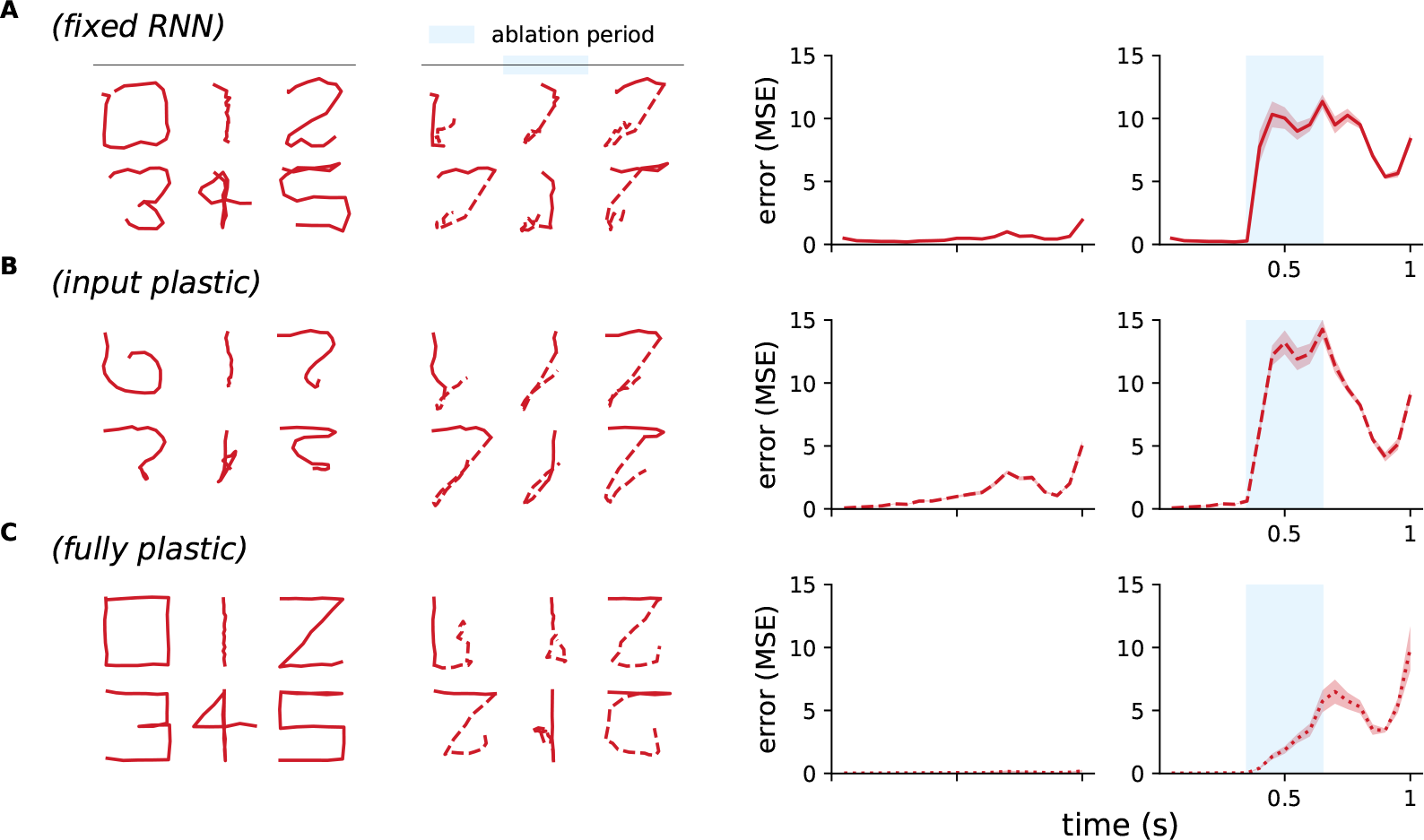
Ablation results for the digit drawing task. **A-C**, Model output (left) and error (right) for digit drawing input under cerebellar ablation for a (A) fixed, (B) input plastic, and (C) fully plastic RNN. Model output shown after cerebellar ablation with dotted lines.

**Figure S10.**
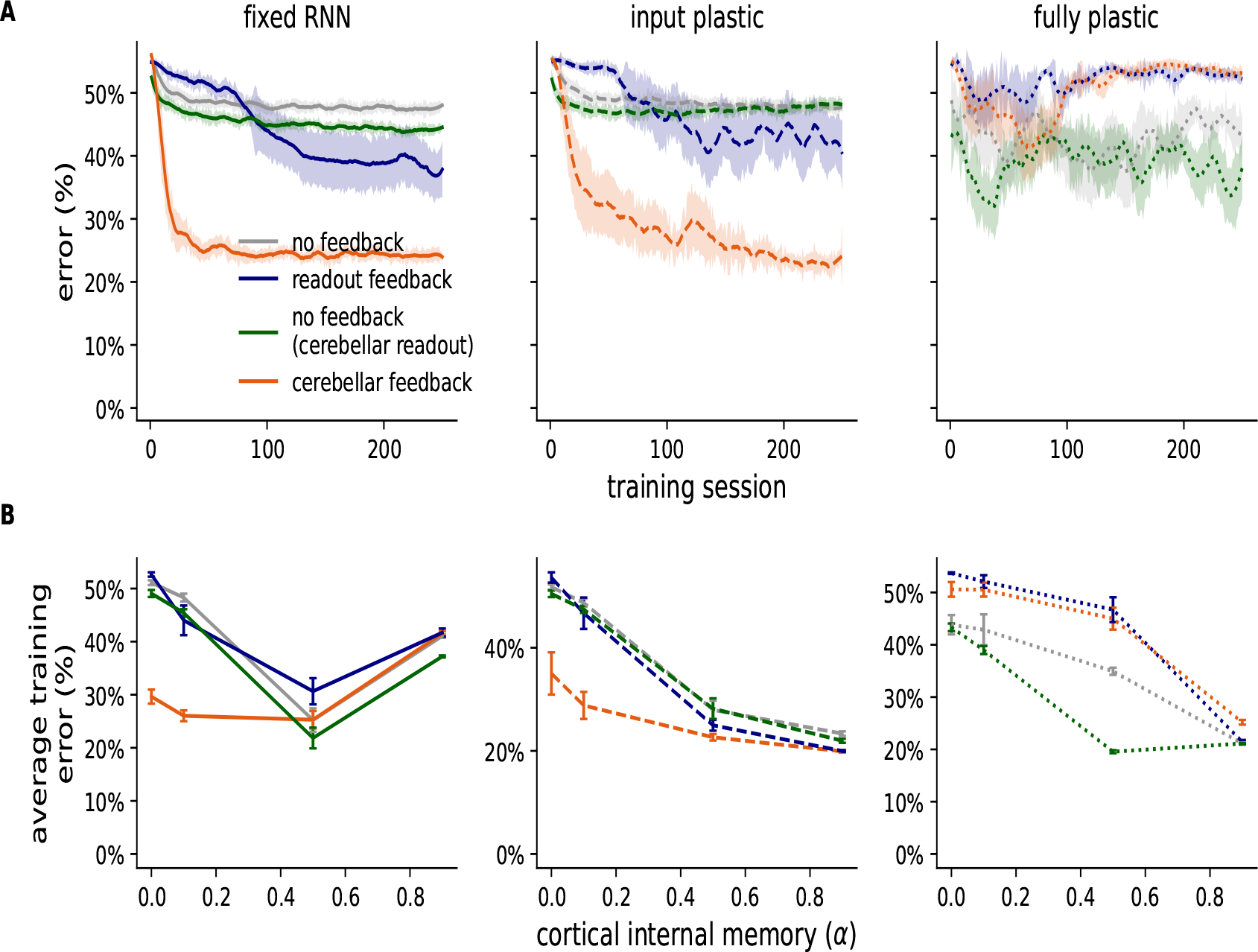
Model learning in the evidence accumulation task. (**A**) Training curves (cortical internal memory *α* = 0.1) for the different models with a fixed RNN (left), input plastic (middle) and fully plastic (right) RNN. Green denotes the model where no feedback is applied to the RNN but the readout network (usually linear) now has the same architecture as the cerebellar network. (**B**) Average error over training across different levels of cortical internal memory *α*.

**Figure S11.**
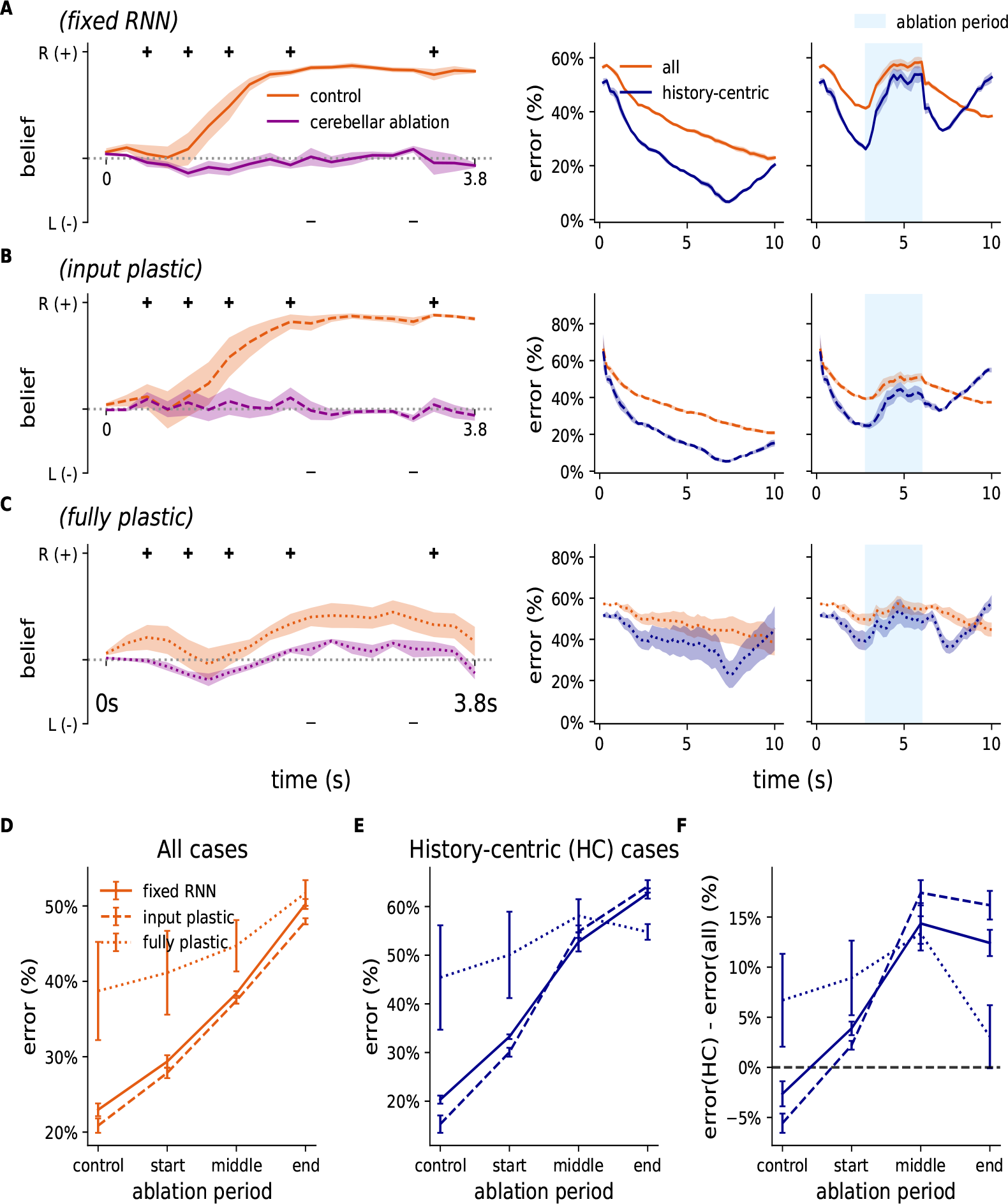
Additional cerebellar ablation results for evidence accumulation task. (**A**-**C**) Model output (left) and error (right) with and without cerebellar ablation (model output shows full cerebellar ablation case) for (A) fixed, (B) input plastic, and (C) fully plastic RNN. (**D**) The error for different ablation periods across these RNN plasticity conditions over all test examples. (**E**) The error for different ablation periods across different RNN plasticity conditions, but only over “history-centric” inputs. (**F**) The difference between D and E.

**Figure S12.**
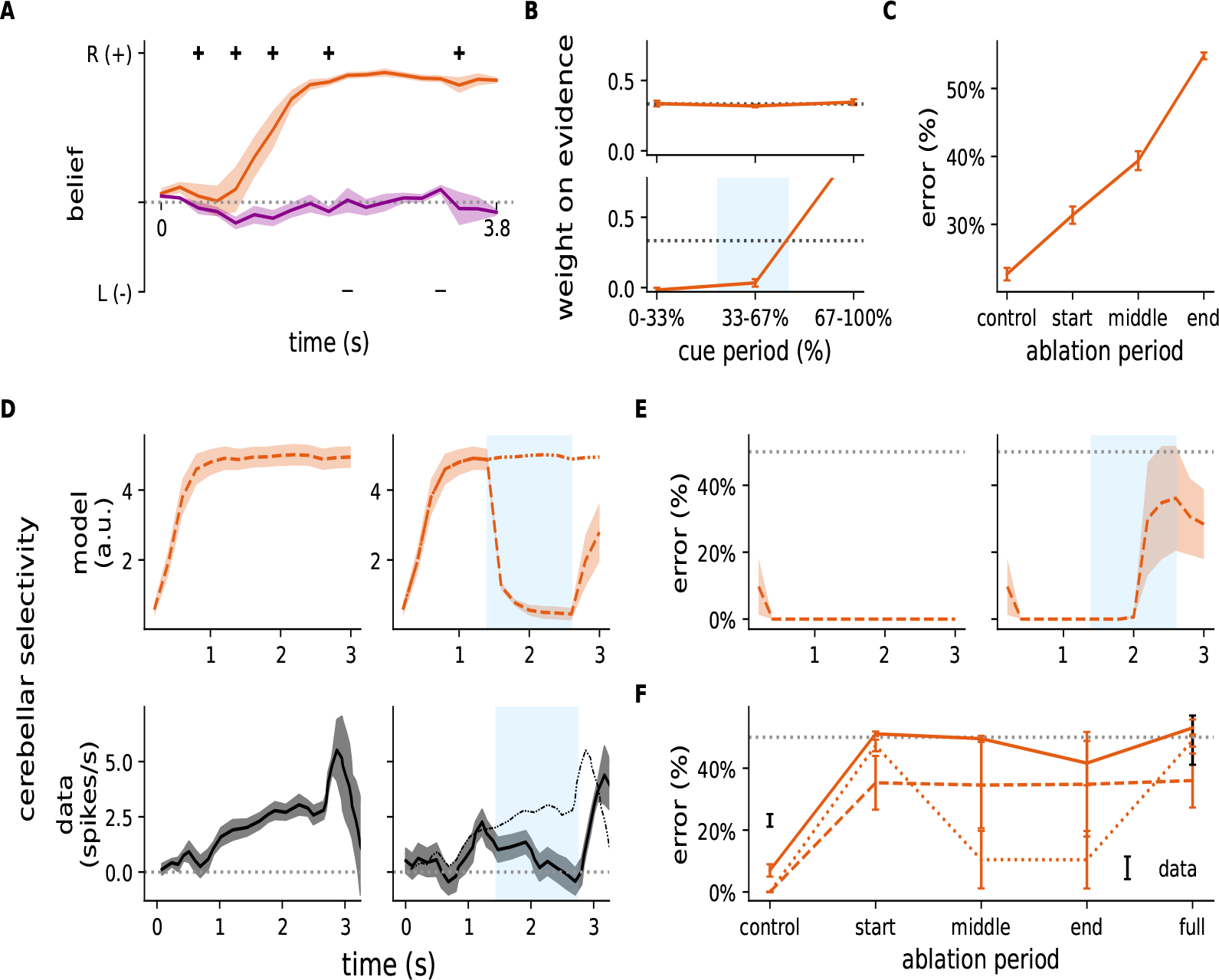
Effect of cortical ablation in working memory tasks. In this analysis 75% of cortical RNN neurons are silenced (after training). (**A**-**C**) Cortical ablation during the evidence accumulation task (fixed RNN). (**A**) Model output without (orange) and without (purple) cortical ablation over the whole task period. (**B**) Normalised regression weights at different periods of input presentation (cue) during control (upper) and cortical ablation (lower; ablation period denoted in blue) conditions. (**C**) Model error under different ablation periods. (**D**-**F**) Delayed association task (input plastic RNN). (**D**) Cue selectivity in the cerebellar network during the delay period without (left) and with cortical ablation (ablation period denoted in blue) conditions for example input in model (upper panels) and experimental data (lower panels) reproduced from Gao et al. ^25^. (**E**) (Cortical) model error during delay period with (left) and without (right) cortical ablation. (**F**) Average error from cortical ablation at different periods during the task delay period and different degrees of plasticity. Experimental data shown in black.

**Figure S13.**
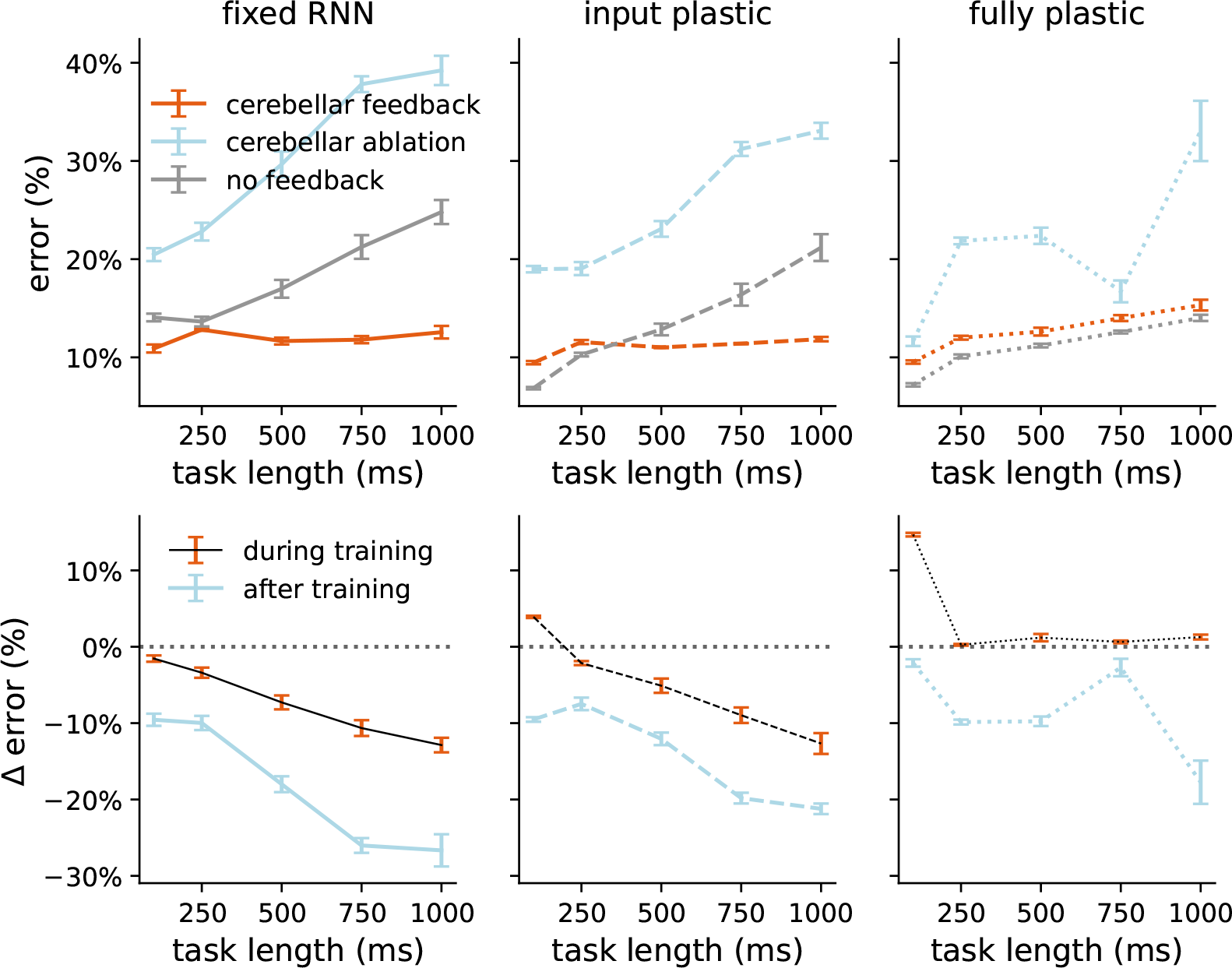
Effect of cerebellar ablation on evidence accumulation task with varying cue durations. (**A**) Test error over different cue durations for models trained with cerebellar feedback (orange), models trained with cerebellar feedback but now subject to cerebellar ablation (light blue), and models trained without cerebellar feedback (grey), with a fixed (left), input plastic (middle) or fully plastic (right) RNN. (**B**) Average change in training error over different cue durations for models with versus without cerebellar component during and after training.

**Figure S14.**
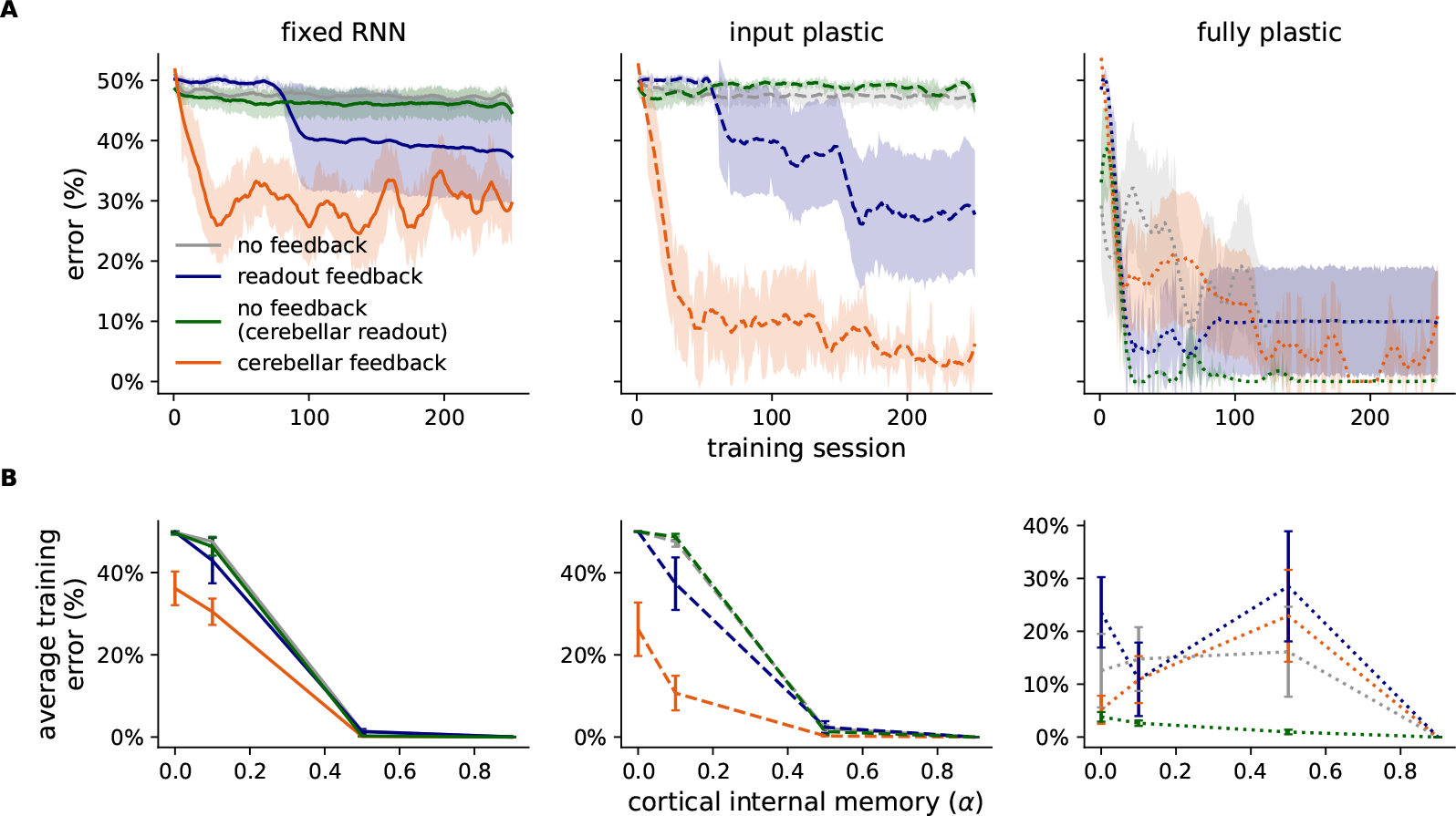
Learning curves in the delayed association task. (**A**) Training curves (cortical internal memory *α* = 0.1) for the different models with a fixed (left), input plastic (middle) and fully plastic (right) RNN. Green denotes the model where no feedback is applied to the RNN but the readout network (usually linear) now has the same architecture as the cerebellar network. (**B**) Average error over training across different cortical internal memory *α*.

**Figure S15.**
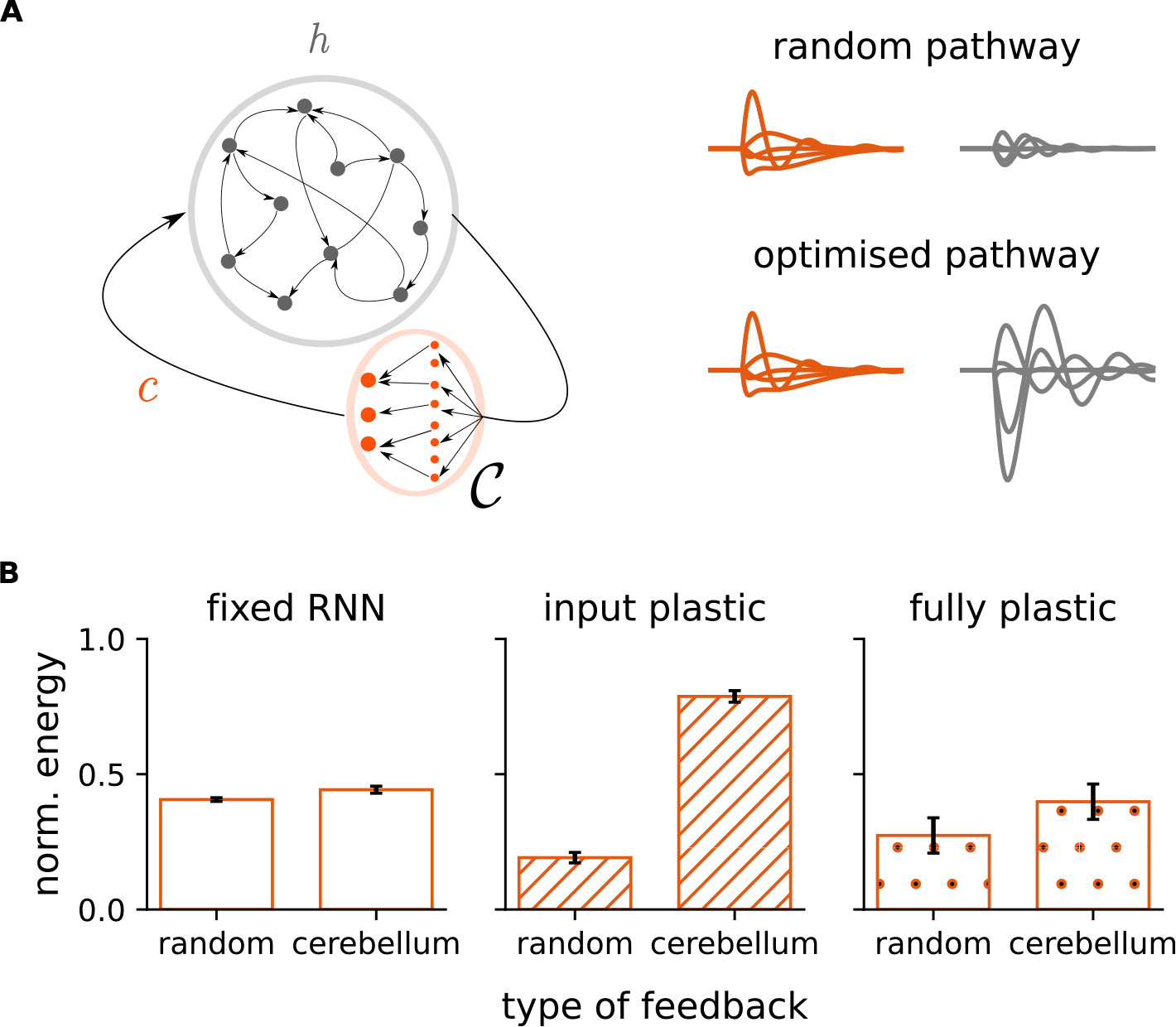
A control-theoretic perspective of the cortico-cerebellar loop. (**A**) Illustrative schematic of cerebellar (orange) and cortical (grey) activities. Depending on the cerebellar-cortical connectivity *W*_*𝒞h*_, the same cerebellar output **c** might suppress (top right) or amplify (bottom right) RNN trajectories. (**B**) The energy (see Methods) generated by random and cerebellar feedback for models trained with varying degrees of plasticity in the delayed association task (Fig. 6). The energy is normalised by the maximum possible energy generated by inputs that achieve the greatest cortical response (see Methods).

**Figure S16.**
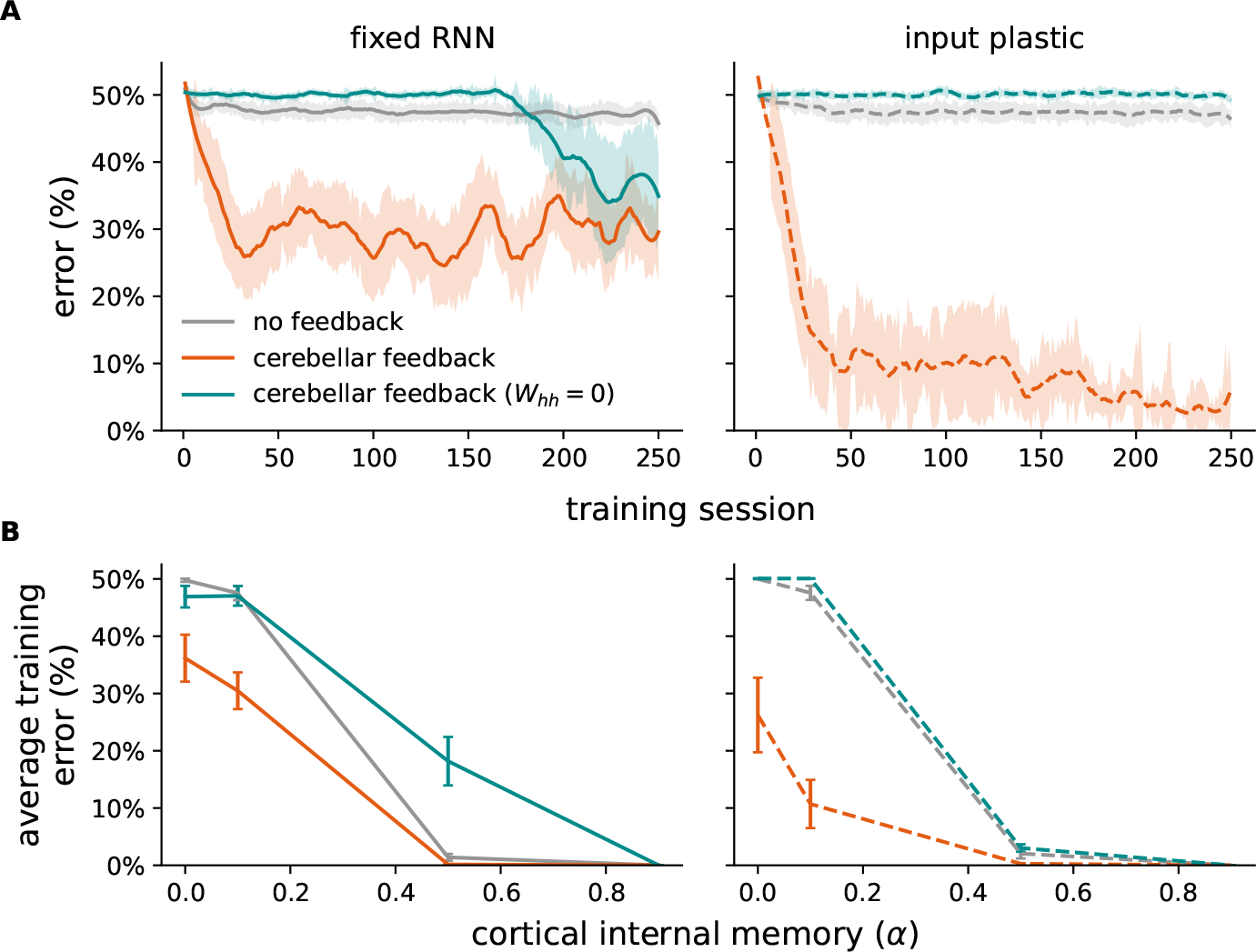
Cerebellar feedback with cortical recurrent connectivity is necessary to learn long-range temporal associations. (**A**) Training curves (cortical internal memory *α* = 0.1) with cerebellar feedback but zero recurrent weights (*W*_*hh*_ = 0) with a fixed (left) and input plastic (right) RNN. (**B**) Average error over training across different cortical internal memory *α*.

**Figure S17.**
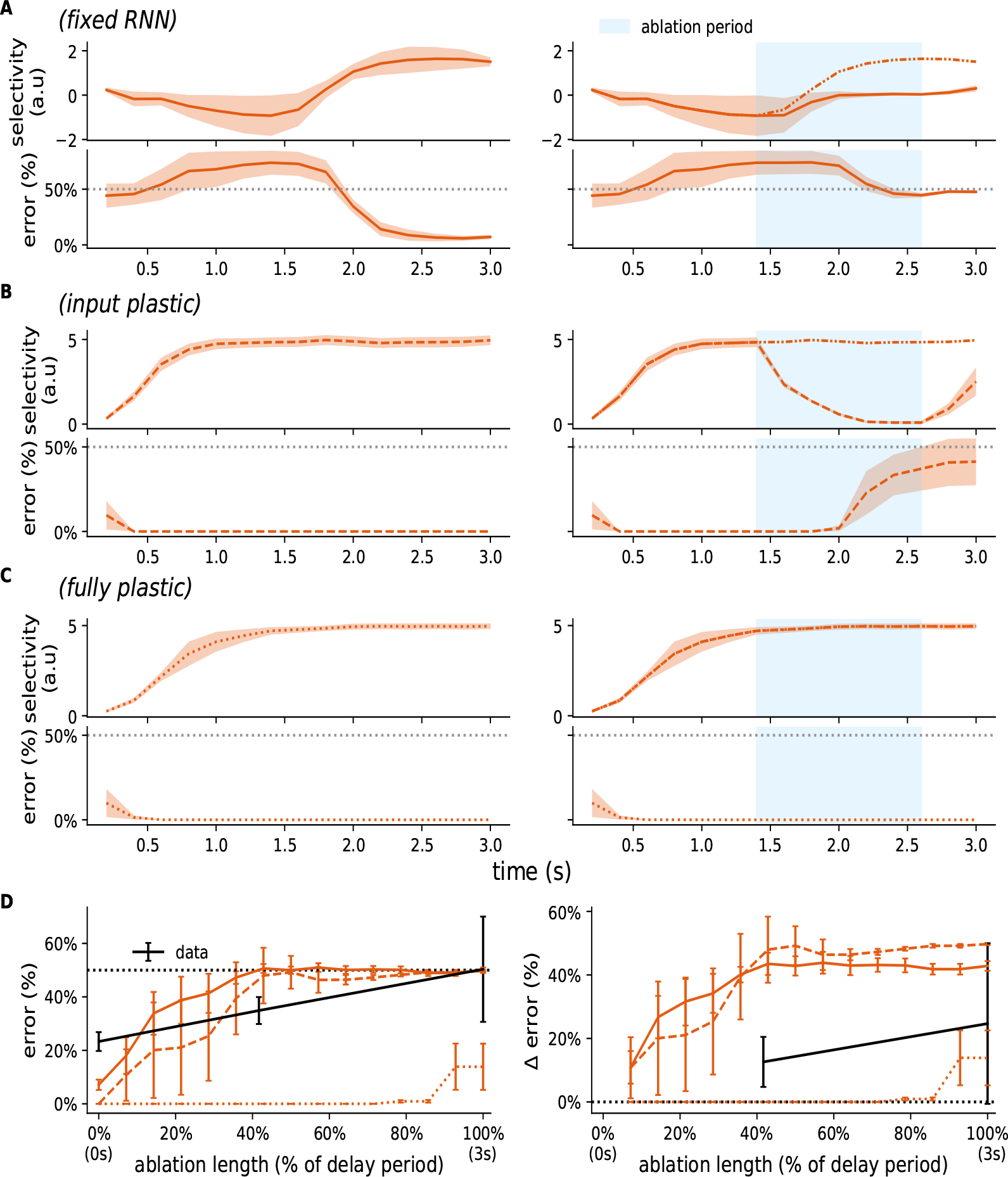
Additional cerebellar ablation results for the delayed association task. (**A**-**C**) Model output (top) and error (bottom) for the delayed association task without (left) and with (right) cerebellar ablation with a (A) fixed, (B) input plastic, and (C) fully plastic RNN. Thin line after ablation shows control model. (**D**) Model error as a function of ablation length (centred around the middle of the delay period). Experimental data reproduced from Gao et al. ^25^. Dotted black line denotes chance.

**Figure S18.**
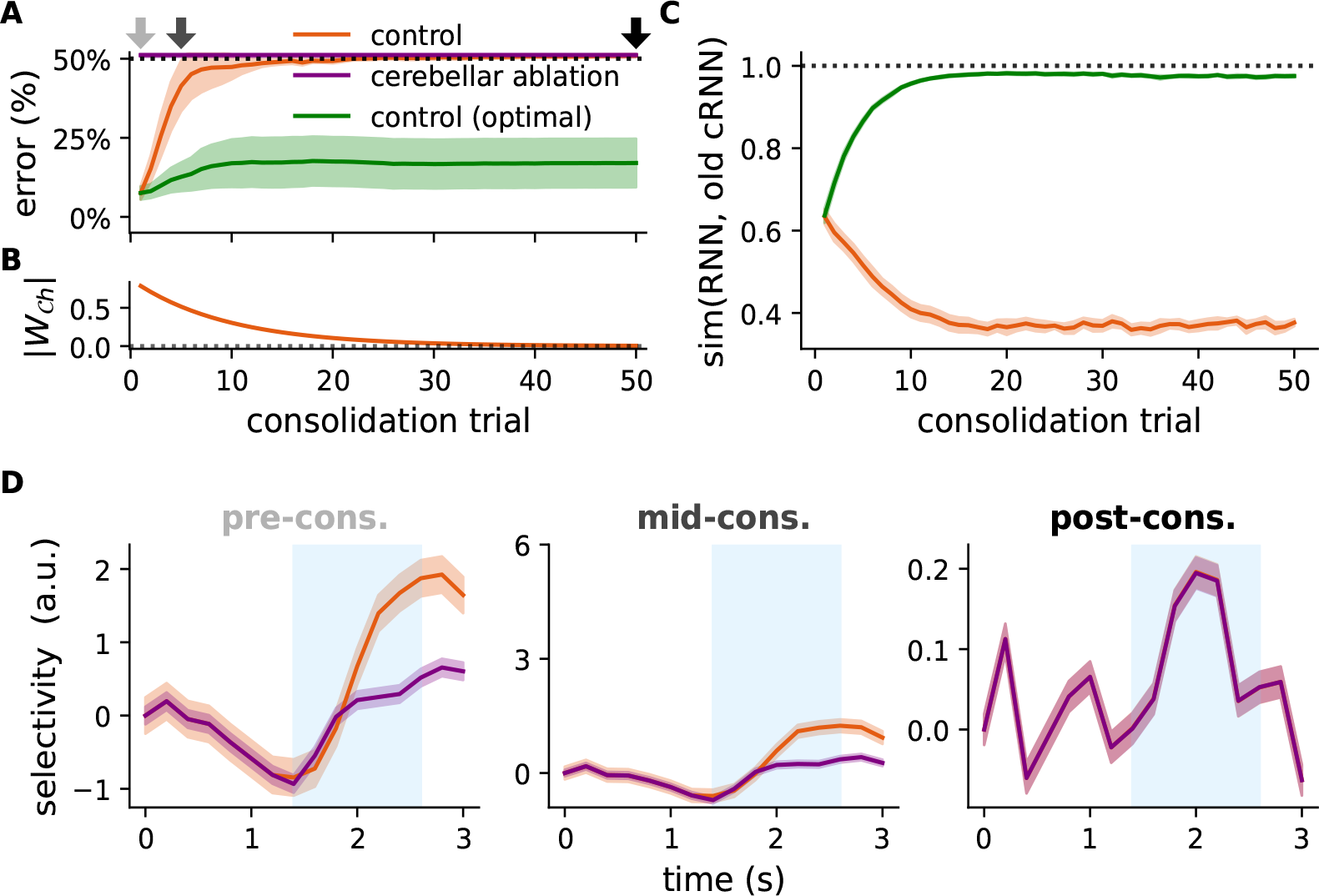
Cerebellar-to-cortical consolidation of the delayed association task with fixed RNN models. (**A**) Accuracy of control and cerebellar ablation conditions (dotted line denotes chance) and the corresponding (**B**) strength of the cerebellar-cortical pathway (*W*_*𝒞h*_) over consolidation. Green denotes control condition with theoretically optimal learning rule. (**C**) Cosine similarity between cortico-cortical input and total cortical input (i.e. cerebellar-cortical and cortico-cortical inputs) pre-consolidation. Similarity of the consolidation model is shown in orange and the optimal consolidation model in green. (**D**) Model selectivity for example (external) input in control and cerebellar ablation conditions at different stages of the consolidation process; colour coded by arrow times in A.

**Figure S19.**
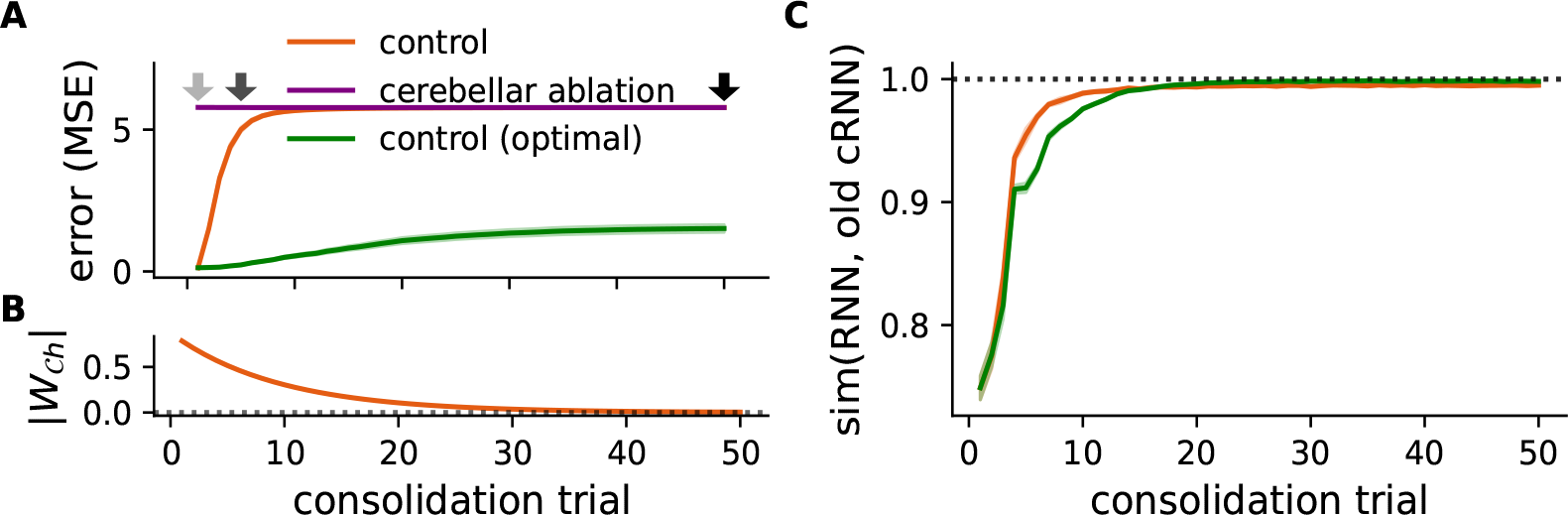
Cerebellar-to-cortical consolidation in linedraw task (fixed RNN). (**A**) Error (mean-squared error) of control and cerebellar ablation conditions and the corresponding (**B**) strength of the cerebellar-cortical pathway (*W*_*𝒞h*_) over consolidation. Green denotes control condition with theoretically optimal learning rule. (**C**) Cosine similarity between cortico-cortical input and total cortical input (i.e. cerebellar-cortical and cortico-cortical inputs) pre-consolidation. Similarity of the consolidation model is shown in orange and the optimal consolidation model in green. Note that even though the similarity between these models is high, their small differences result in significant changes in the overall trajectory of cortico-cerebellar activity, resulting in poor final performance.

